# Redistribution of codon-optimality effects: measurement strategy alters the division of labor between translation and mRNA decay

**DOI:** 10.64898/2026.05.21.726845

**Authors:** Sayanur Rahaman, Samuel Mondal, Colin E. Delaney, Misaal Bedi, Sandrine Wallerich, Chandraday Prodhan, Vincent Jaquet, Attila Becskei

## Abstract

Codon optimality promotes efficient translation and, as recent research has shown, also extends mRNA lifetimes. However, how control is distributed between translation and mRNA degradation remains unclear. We show that this relative impact depends strongly on the measurement approach. Using fluorescent protein reporters can underestimate codon-optimality–dependent increases in translation efficiency. Conversely, analyses based on poly(A)-selected RNA overestimate the impact on translation, because stable transcripts undergoing poly(A) shortening are often inefficiently captured, leading to skewed protein-to-mRNA ratios. This technical bias is not offset by the marginal decline in ribosomal association observed as mRNAs age. Estimates based on total RNA measurements redistribute some of the control attributed to translation to mRNA stability, making the contributions comparable for mRNAs with shorter coding sequences. For longer mRNAs, codon optimality increasingly controls elongation speed, with a greater effect on translation efficiency than on degradation. These insights highlight the importance of measurement strategy for accurately quantifying the determinants of mRNA stability and protein synthesis.

## INTRODUCTION

Post-transcriptional processes, in particular the translation and degradation of mRNAs, play a major role in shaping gene expression. The rates of translation and mRNA decay are shaped by multiple factors, with codon optimality emerging as a key determinant. Most amino acids are encoded by several synonymous codons, but these codons are not used equally across genes. It has long been observed that some genes exhibit a strong codon bias, preferentially using certain synonymous codons while avoiding others, whereas other genes show little or no such bias^1,2^. The observation that highly expressed genes—particularly ribosomal protein genes—favor certain codons led to the introduction of the codon adaptation index (CAI) as a quantitative measure of codon bias. These genes tend to rely on codons recognized by abundant tRNAs, whose availability is largely determined by their genomic copy number^3^. This relationship to translation has long been recognized: codons read by abundant tRNAs generally promote faster and more efficient elongation.

More recent work has shown that codon optimality also plays a major role in determining mRNA half-lives. This connection was slower to emerge, partly because the mechanistic link is less direct. In eukaryotic organisms ranging from yeast to human, codon-dependent elongation speed influences decay through the Ccr4–Not complex, which recognizes ribosomes paused at non-optimal codons while awaiting tRNAs^4–6^. However, the extent to which codon usage influences mRNA stability has often appeared inconsistent across studies. A key reason for this discrepancy is the strong dependence of measured half-lives on experimental design—most notably the methods of RNA isolation and half-life determination, which can themselves perturb the mechanisms regulating mRNA stability^7,8^. Whereas most recent studies reported that optimal codons prolong mRNA half-lives, many earlier datasets failed to show a clear correlation, and in a few cases even displayed opposite trends depending on experimental design and RNA isolation^9^.

Since translation efficiency (TE) depends on both protein and RNA measures, we hypothesized that methodological biases affecting mRNA half-life estimation would also influence the assessment of TE. This analysis revealed that both translation and degradation display very similar codon dependence. At the same time, we find that measurements of mRNA stability and translation efficiency (TE) are strongly affected by whether total RNA or poly(A)-selected RNA is used. Because poly(A)^+^ isolation with oligo(dT) beads is widely applied in transcriptomic studies, it is essential to understand how these methodological choices shape the apparent relationships between translation and decay.

Poly(A)-tail processing plays an important role in determining which populations of poly(A)+ mRNAs are isolated, which in turn affects measured mRNA turnover and translation. After transcription, mRNAs are capped at the 5′ end of the 5′UTR and polyadenylated at the 3′UTR^10–12^. The cap is required for translation initiation, and once exported to the cytoplasm, the mRNA is rapidly engaged by ribosomes. At the same time, its poly(A) tail is gradually shortened by deadenylases^13^. Once the tail is eliminated or is reduced to a critical length, the mRNA is decapped, which permits degradation by Xrn1 in the 5′–3′ direction^11^. mRNAs lacking a poly(A) tail can still be translated^14^. As with the relationship between mRNA stability and codon optimality, the relationship between tail length and mRNA stability has historically been subject to significant ambiguity. For instance, tail length and half-life were reported to correlate either negatively or positively depending on the source of half-life measurements, with Spearman correlation coefficients ranging from −0.44 to 0.23^15^.

Here, we built a mathematical model to describe how poly(A)-tail dynamics and RNA isolation strategies influence apparent mRNA half-lives. We then used this framework to examine the quantitative partitioning of codon-optimality-dependent control between translation and degradation, a relationship that has not yet been systematically analyzed. We show that the apparent balance between translational and decay control depends strongly on measurement strategy. Moreover, elongation-dependent effects on translation become increasingly dominant for mRNAs with longer coding sequences.

## RESULTS

### Unified Datasets Reveal Strong Correlation Between Codon-Mediated Stability and Translation

To investigate the interplay between mRNA degradation and translation in the model organism budding yeast, we integrated multiple genome-wide datasets to enable more precise and comprehensive comparisons. For mRNA half-lives, we relied on a previously unified dataset^16^. In this work, we define Translation Efficiency (TE) as the protein synthesis rate constant (protein molecules produced per mRNA per unit time, Equations (10), (11)). This definition captures the combined effects of initiation, elongation, and termination, distinguishing it from ribosome density measured by ribosome profiling (Ribo-seq). While Ribo-seq provides high-resolution snapshots of ribosome positions, it does not directly capture elongation speed; therefore, a higher ribosome density does not necessarily equate to a higher protein synthesis rate if elongation is slow.

To analyze TE, we gathered three datasets (Fig. 1A, Table S1) based on stable isotope labeling with amino acids and mass spectrometry (SILAC-MS) combined with mRNA quantification to determine protein synthesis rates^17–19^. The TE values reported in these studies showed weak to moderate correlations with one another (Fig. S1A). Their interquartile ratio ranged from 6 to 9 (Fig. 1B).

**Figure 1.**
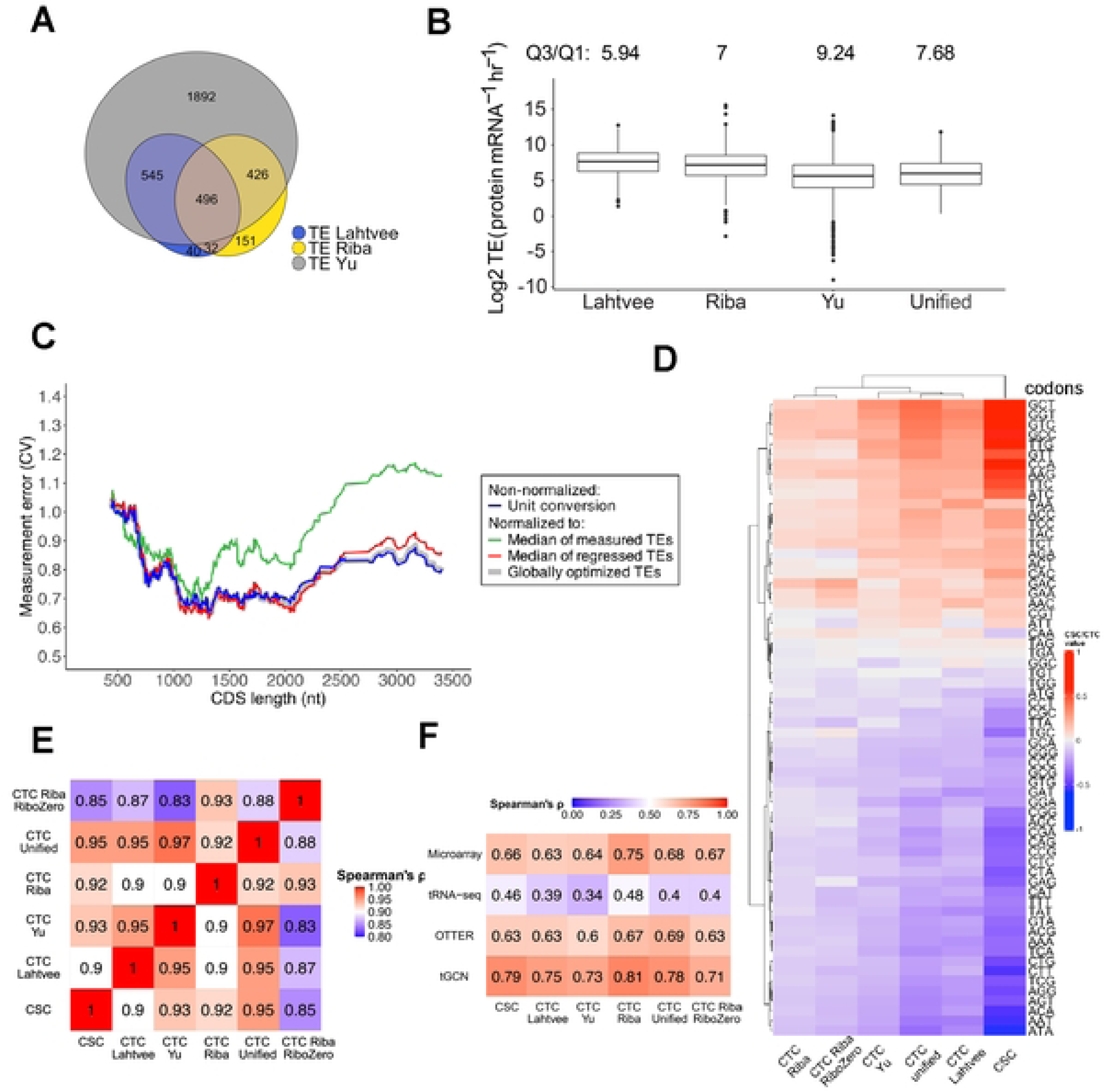
Comparison of mRNA codon stabilization (CSC) and codon translation (CTC) coefficients in yeast. **(A)** Venn diagram illustrating the overlap of genes across TE datasets (Lahtvee, Riba, Yu). A total of 3,582 mRNAs are included, of which 496 are common to all datasets. (**B)** Boxplots of log₂ TE values for the individual datasets and the unified dataset. The interquartile ratio (Q3/Q1) is shown above each plot; Q1 and Q3 represent the 25th and 75th percentiles, respectively. (**C)** Measurement error, expressed as the coefficient of variation (CV), calculated for the 496 genes present in all datasets. Rolling mean CV is plotted against CDS length for non-normalized (unit converted) TE values and for three normalization strategies (median-normalized, regression-based median, globally optimized scaling factors). (**D)** Hierarchically clustered heatmap of CSC and CTC values. CSC values were derived from half-life data for 82 genes; CTC values were derived from TE datasets: Lahtvee (1,113 genes), Yu (3,363 genes), Riba/RiboZero (1,106 genes), and the unified dataset (3,278 genes). (**E)** Spearman correlation matrix of CSC and CTC values across datasets, calculated from codon frequency correlations with either half-life or TE for all 64 codons. For clarity, only values ≥0.8 are shown. (**F)** Correlation of CSC and CTC values with independent measures of tRNA abundance or activity (microarray, tRNA-seq, oligonucleotide-directed three-prime terminal extension of RNA (OTTER), tRNA gene copy number (tGCN)). Correlations were calculated for 41 tRNA isoacceptors; microarray data lack values for two isoacceptors (anticodons CTG and CTC).

To unify the datasets, we converted all TE values to a common unit and calculated the mean TE for each gene excluding those with large discrepancies across datasets. As a result, we obtained a unified dataset with TE data for 3278 mRNAs. To assess the quality of the unified datasets, we calculated the variability of TE measurements across the three datasets and its dependence on coding sequence (CDS) length. TEs showed the lowest variability around the most common CDS length, between 1000 and 2000 nucleotides (nts) (Fig. 1C) and the variability is higher for short (less than 1000 nts) and long mRNAs (more than 2000 nts). Similar trends were observed for mRNA half-lives^16^. Interestingly, alternative forms of normalization, such as median based normalization prior to averaging failed to further reduce the variability in the unified TE (Figs. 1C and S1B), suggesting that this simple form of unification reaches a high quality.

mRNAs with very high TE predominantly encode the highly expressed ribosomal proteins or glycolytic enzymes (Fig. S1C). To examine the relationship between TE and codon optimality, we applied an approach analogous to that used for mRNA stability. Codon effects on mRNA half-lives have been commonly quantified via the codon stability coefficient (CSC), which is defined as the correlation between codon frequency and mRNA half-life^4^. Similarly, we defined a codon translation coefficient (CTC) to assess how codons influence TE (Fig. 1D). CTCs from each dataset correlated strongly with the CSCs (typically ρ ≈ 0.9; Fig. 1E), indicating that codons promoting efficient translation also tend to stabilize mRNAs. Notably, the unified dataset’s CTCs showed an even stronger correlation with CSCs (ρ = 0.95), reinforcing the robustness and reliability of the unification approach.

CTC and CSC characterize codons from the perspectives of translational efficiency and mRNA stability, respectively, thereby revealing how individual codons enhance these processes. Consistently, we observed a strong correlation between both parameters and the CAI (Fig. S1E), which reflects biased codon usage. Since codons exert their effects primarily through decoding by tRNAs, we compared CTC and CSC with parameters that characterize tRNAs and their interactions with codons rather than properties of the codons themselves. The primary determinant of tRNA efficacy is their intracellular abundance, which is directly proportional to the copy number of tRNA genes in the genome^20^. Both CSC and CTC show strong and comparable correlations with tRNA copy numbers and abundances, particularly when measured by microarray rather than RNA-seq (Fig. 1F). The OTTER method (based on terminal RNA extension) also yields high correlation^20^. Notably, incorporating wobble-pairing effects—by extending the analysis from 41 directly measured isoacceptors to all 61 sense codons—did not further improve these correlations (Figs. 1F, S1D). These findings indicate that both mRNA stability and translation benefit from the same core principle: codons recognized by abundant tRNAs promote efficient translation and stabilize the transcript, whereas codons read by low-abundance tRNAs yield weakly translated, short-lived mRNAs.

### Effects of mRNA isolation and detection methods on mRNA half-life estimates

In the three datasets described above, TE was calculated as the ratio of protein production rate to mRNA abundance. For mRNA quantification, two datasets used poly(A)+ RNA selection—the standard positive-selection approach that enriches mRNAs with poly(A) tails using oligo(dT) beads. In the third study, two TE estimates were provided: one based on poly(A)+ selection (Riba dataset), as above, and another based on rRNA depletion (Riba RiboZero dataset), a negative-selection method that enriches mRNAs by removing rRNAs. Because rRNA depletion removes ribosomal RNA without selecting for poly(A) tails, the resulting mRNA population is effectively equivalent to total mRNA.

The importance of the RNA isolation method is underscored by previous findings showing that it can influence measured mRNA half-lives. In particular, poly(A)+ selection has been reported to abolish the correlation between CSC and tRNA gene copy number or abundance, at least when half-lives are determined with transcriptional inhibition^4,9^. This reduction has been attributed to the selective stabilization of certain mRNAs upon poly(A) tail shortening^4^, as deadenylation typically precedes decapping^21^.

We next calculated CTCs using both Riba datasets to assess the effect of RNA isolation on TE. Since poly(A)+ selection is known to bias half-life measurements, we expected that rRNA depletion—which captures the total mRNA population—would provide a more accurate reflection of codon-driven translation. However, contrary to this expectation, the CTCs derived from rRNA depletion correlated less with tRNA gene copy number than those derived from poly(A)+ RNA (Fig. 1F, r_S_ = 0.81 vs 0.71; p = 0.0086, n = 41; Meng-test). Similarly, the correlation of these CTCs with the CSC was significantly different (r_S_ = 0.92 vs 0.85; p = 0.00055, n = 64; Meng-test^22^).

Because correlations of codon optimality with half-life and TE responded differently to RNA isolation method, and given the widespread use of poly(A)+ selection in gene expression studies, we sought to validate these observations. We first re-examined whether RNA isolation affects mRNA half-lives, since methodological differences can significantly impact interpretations and conclusions^9^. While the above differences in half-life were observed using transcriptional inhibition, we asked whether similar effects would be observed with the independent gene control method^7^.

To this end, we used a *tet-*promoter to shut off transcription of selected mRNAs, including both stable and unstable transcripts. After adding doxycycline, samples were collected at the indicated time points, and RNA was isolated. mRNA levels were quantified either from total RNA samples or from poly(A)^+^ RNA isolated using pull-down with oligo(dT) coated beads, a commonly used mRNA enrichment strategy in RNA-seq workflows (Fig. 2A). For the relatively stable *PRO3* mRNA, the half-life was shorter when measured from poly(A)+ RNA (54% of the half-life determined from total RNA; P = 0.001, t-test on fitted half-lives with propagated SEs). In contrast, the unstable *DUT1* mRNA showed no significant difference (poly(A)+ RNA half-life: 89% of that measured in total RNA; P = 0.47). These results recapitulate the trend seen with the half-lives obtained with transcriptional inhibition^4^.

**Figure 2.**
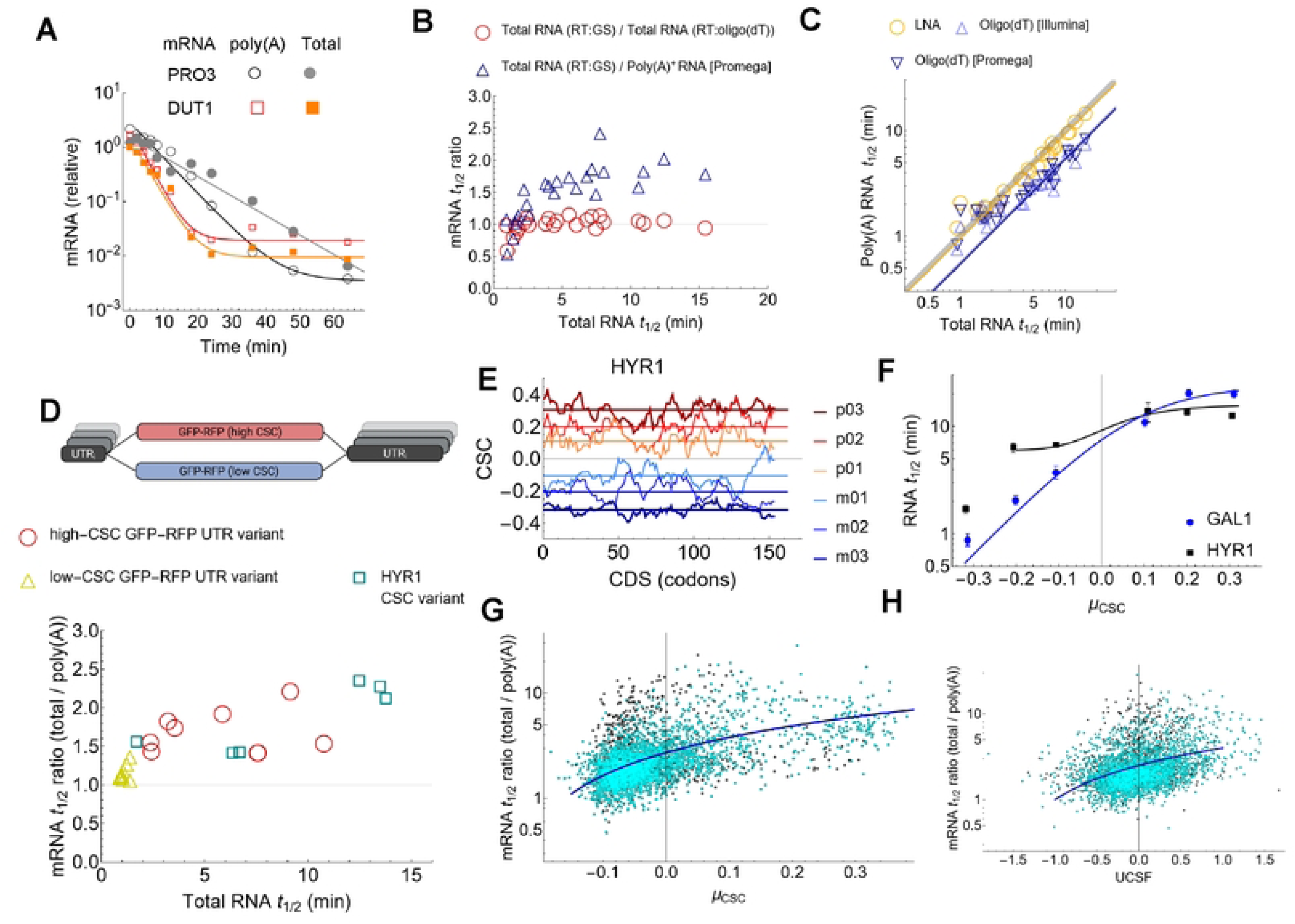
**Systematic differences in mRNA half-lives determined from using total RNA and poly(A)^+^ selected RNA and their dependence on codon optimality**. **(A)** Decay of mRNAs measured (relative to housekeeping genes) at the indicated time points following doxycycline addition. Reverse transcription (RT) was performed with gene specific primers on either total RNA or poly(A)^+^ mRNA isolated with oligo(dT) beads (Promega). Half-lives were determined by nonlinear regression for the stable *PRO3* (*YER023W*), 8.4 ± 1.1 min (total) and 4.6 ± 0.3 min (poly(A)^+^); and the unstable *DUT1* (*YBR252W*), 2.7 ± 0.3 min (total) and 2.4 ± 0.3 min (poly(A)^+^). **(B)** Half-lives determined by RT-qPCR using gene specific (GS) primers. Results for total RNAs were compared to those obtained by oligo(dT)-primed reverse transcription (total RNA, RT: oligo(dT)) or after poly(A) mRNA pulldown with oligo(dT) (Promega; poly(A) isolation, RT: gene specific). Both half-life ratios correlated (*r_s_*) positively with the total mRNA half-lives but only the pull-down half-life ratios were significantly different (RT with oligo(dT): *r_s_* = 0.47, P = 0.02; signed-rank test P = 0.064; pull-down with oligo(dT): rs = 0.86, P = 7.2 10^-8^; signed-rank test P = 0.0001). **(C)** Comparison of the half-lives of RNAs quantified with total RNA and poly(A)+ RNA isolated with oligo(dT) or LNA oligo(T) beads. The isolation with LNA yields half-lives that do not differ significantly from those from total RNA (P = 0.31, Signed Rank Test; slope = 0.94), whereas the poly(A)-RNA isolated with oligo(dT) differs significantly (P = 10^-4^, slope = 0.53 for method TruSeq (Illumina), and P =10^-4^, slope = 0.56 for Promega). The thick gray line denotes the identity line. **(D)** Scheme of the GFP-RFP high and low-CSC mRNAs (top panel). Half-life ratios of the UTR variants of GFP-RFP and CSC-recoded *HYR1* mRNAs (bottom panel) as a function of half-lives measured from total RNA. Across all constructs, the correlation is significantly positive (r_s_ = 0.81, P =2×10^-7^). However, this correlation is not significant when analyzing the high-CSC UTR series (r_s_ = 0, P =0.96) or the low-CSC UTR series (r_s_ = 0.13, P =0.74) independently. **(E)** Recoding of codon optimality in the coding sequence of the *HYR1* mRNA. The lines and the curves represent the global and moving averages (with 10 codon length windows) of the recoded constructs. The µ_CSC_ values for the recoded variants are m03 =-0.32, m02 = −0.21, m01 = −0.11 and p01 = 0.11, p02 = 0.20 and p03=0.31. **(F)** Half-lives of the recoded *GAL1* and *HYR1* mRNAs. Error bars denote SEM (*n* = 3 replicates). **(G)** Dependence of the total/poly(A) mRNA half-life ratio on µ_CSC_. Grey dots show all data points from the Presnyak dataset, and cyan dots indicate the outlier-filtered dataset. The dark blue line represents linear regression fit (y = 2.72 + 10.77x). Correlation coefficients: r_s_ (all) = 0.44, r_s_ (outlier-filtered) = 0.52. See Fig. S2C, D for further details. **(H**) The half-life ratio as a function of UCSF; y = 2.51 + 1.51x, r_s_(all) = 0.320, r_s_(outlier-filtered) = 0.341.

Extending this analysis to a broader set of mRNAs, we measured half-lives spanning from 0.5 to 15 min using total RNA with gene-specific primers for reverse transcription (RT:GS). We also measured the corresponding half-lives from poly(A)+ mRNA isolated with oligo(dT) beads. The ratios of these measurements were plotted as a function of total mRNA half-life. For unstable mRNAs, the ratio was close to one, indicating negligible isolation bias, whereas for stable mRNAs the ratio was typically between 1.5 and 2 (Fig. 2B; Poly(A)^+^). The protocol used for these experiments (Fig. 2C, Promega) yielded similar results to that used in RNA-seq (Fig. 2C; Illumina).

The above results confirmed that RNA isolation affects the estimated half-lives of stable mRNAs. We next asked how codon optimality and non-codonic influences, primarily mediated by UTRs, contribute to the observed differences in half-lives between poly(A)+ and total RNA. To address this, we analyzed two sets of constructs in which codon usage and UTR sequences were systematically varied.

We first analyzed synthetic constructs encoding a GFP–RFP fusion protein, which has a CDS length (1.4 kb) close to transcriptome average in yeast. Each construct featured either optimal (high CSC) or non-optimal (low CSC) codons (Fig. 2D). Both CDSs were flanked by various UTRs, which can modulate the stability of mRNAs. These were type II UTRs which can modulate half-life even when the length of the CDS exceeds 400 nts. This distinguishes them from type I UTRs, whose half-life modulating effect is restricted to short CDSs. Most type II UTRs used here had been shown previously to destabilize mRNAs with optimal codons^16^; thus, they effectively decouple mRNA stability from translation. The mean half-life ratio of total to poly(A)-selected RNA was considerably higher for optimal than for non-optimal CDSs (1.69 vs. 1.18, P-value for t-test: P = 0.001) whereas no clear UTR-specific effects were observed (Fig. 2D).

Our earlier work introduced the ultra-codonic stabilization factor (UCSF) to capture non-codonic, primarily UTR-mediated contributions to mRNA stability^16^. UCSF complements the average codon stability coefficient (µ_CSC_), where µ_CSC_ represents the average CSC value across all codons within the coding sequence (CDS). UTRs with negative UCSF tend to destabilize transcripts, whereas the effect of UTRs with positive UCSF is less clear^16^. To explore stabilizing UTR effects, we examined an mRNA with a highly positive UCSF (1.39), *HYR1*, making it a candidate for detecting UTR-mediated stabilization. Furthermore, it has a short CDS (492 nt), extending our analysis to shorter CDSs. For comparison, we included *GAL1*, which has a relatively neutral UCSF (–0.27).

We generated six variants of each gene by systematically recoding codon optimality, varying µ_CSC_ from –0.3 to 0.3 (Fig. 2E). For both mRNAs, the half-lives plateaued between 10 and 20 min within the positive µ_CSC_ range (Fig. 2F). In the negative range, we focused on µ_CSC_ = –0.2, which represents the lower limit of codon optimality observed among endogenous transcripts (Fig. 2G). At this value of µ_CSC_, the *HYR1* half-life was approximately threefold longer than that of *GAL1*, and declined only at µ_CSC_ = –0.3—a codon composition not observed naturally. Thus, the *HYR1* UTR can stabilize its CDS across the genomically relevant range of µ_CSC_. Together, these findings indicate that UTRs with highly positive UCSF can stabilize mRNAs with suboptimal codon usage, identifying a novel variant of stabilizing Type II UTRs to complement previously known destabilizing variants.

Across all *HYR1* constructs, the ratio of total-to-poly(A)+ half-lives correlated strongly with half-lives measured from total RNA (Pearson correlation, *r* = 0.87), confirming that codon usage continues to influence mRNA stability even in shorter CDS contexts with stabilizing UTRs (Fig. 2D). Furthermore, their ratios were similar to that found for the high-CSC GFP-RFP mRNAs (P = 0.58, t-test), and significantly higher than that of the low-CSC GFP-RFP mRNAs (P = 0.04). This confirms that UTRs with a highly positive UCSF can effectively phenocopy the stabilizing effect of optimal codons across different gene contexts.

In summary, our findings confirmed the effect of codon optimality and stabilizing UTRs. Similar to endogenous mRNAs, the ratios were close to one when total mRNA half-lives were below ∼2 minutes (Fig. 2B). This indicates that the divergence between poly(A)+ and total half-lives emerges above the median half-life of the transcriptome (2 min).

With the limited number of synthetic constructs tested, the dependence of the total-to-poly(A)+ half-life ratio on UTRs did not reach significance (Fig. 2D). We therefore extended the analysis to the transcriptome level, using a outlier-filtered version of the half-life dataset from the transcriptional inhibition (TI, Presnyak) study (Methods)^4^. The poly(A)+ to total half-life ratios obtained with the multiplexed gene control correlated positively with the Presnyak dataset (r_s_ =0.53), a correlation that strengthened after outlier filtering (r_s_=0.75) (Fig. S2A).

The transcriptome-wide half-life ratio correlated positively with µ_CSC_ and UCSF (Figs. 2G, 2H). Notably, the correlation was significantly stronger with µ_CSC_ than with UCSF (r_S_ = 0.52 vs 0.34; Meng’s test for dependent correlations: p = 1.7×10^-18^)^22^. Given the overall correlation between total half-life and the poly(A)+/total ratio (r_S_ = 0.59, Fig. S2D), we conclude that codon optimality is the dominant factor underlying differences between poly(A)+ and total half-lives, with UTR-mediated effects playing a smaller but detectable role.

### Oligo(dT) Beads Impose an (A)₂₀ Threshold

These results indicate that RNA isolation affects the measured half-lives of stable mRNAs, particularly those with optimal codons and, to a lesser extent, those with stabilizing UTRs. Given the widespread impact of isolation and detection methods, we next examined the methodological basis of these differences.

Interestingly, when oligo(dT) was used solely as a primer for reverse transcription on total RNA (RT: oligo(dT)), the half-life ratio was close to one for all mRNAs (Fig. 2B). This suggests that the annealing to the beads is a critical factor. We therefore asked whether poly(A)+ capture efficiency could be improved. To this end, we replaced the oligo(dT) in the beads with locked nucleic acid (LNA)-modified oligonucleotides, which contain LNA(T) and exhibit higher affinity for poly(A) tails^23^. The half-lives measured from LNA-selected poly(A)+ mRNA closely resembled those obtained from total RNA, contrasting with the significant differences seen when oligo(dT) beads were used (Fig. 2C). Both methods removed 18S ribosomal RNA with comparable efficiency (depletion percentage: 98.1 ± 0.1% with oligo(dT), 98.3 ± 0.2% with LNA). These results indicate that LNA-oligo(T) beads efficiently capture poly(A)+ RNA without compromising RNA purity, including transcripts with short tails, and yield half-life measurements similar to total RNA or oligo(dT)-priming.

Since reverse transcription and bead-based selection both rely on oligo(dT), the discrepancy between them, together with the results obtained using LNA-modified oligonucleotides, suggests that binding stringency underlies these differences. To characterize this, we determined the poly(A) tail-length threshold captured by oligo(dT). We therefore generated barcoded DNA constructs encoding CDS and UTR sequences with defined poly(A) tail lengths and transcribed them in vitro. Subsequently, mRNAs were reverse transcribed using oligo(dT) primers, and RNA abundance was quantified by qPCR via barcode-specific amplification. Despite some variability, the detected RNA levels were largely proportional to the input RNA, and even poly(A)-negative mRNAs were efficiently primed and detected (Fig. 3A). These results confirm our previous observations that oligo(dT)-based priming confers little or no specificity. In contrast, mRNA recovery declined sharply upon isolation with oligo(dT) beads when poly(A) tail length was reduced below ∼20 nucleotides.

**Figure 3.**
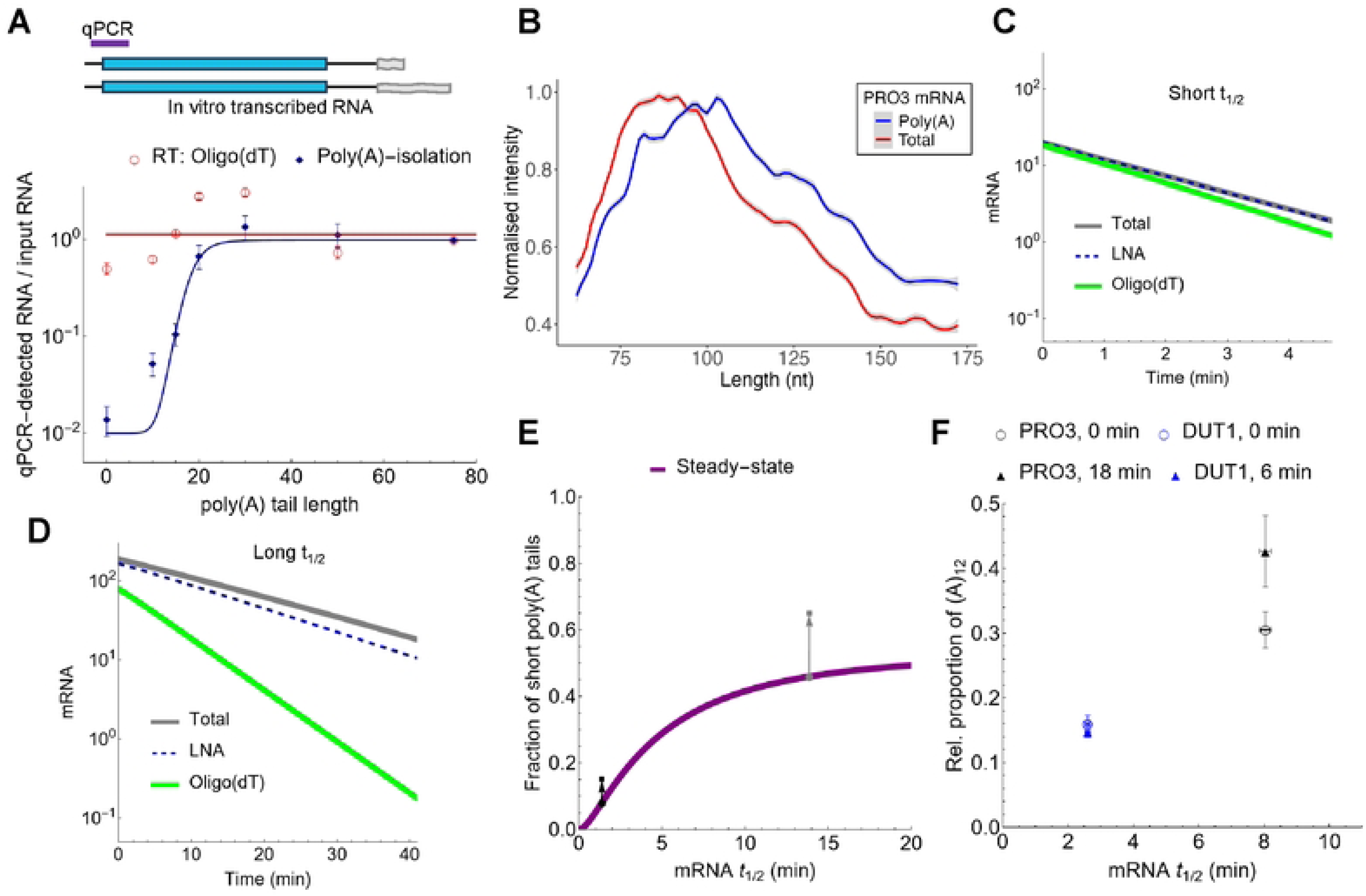
Threshold poly(A) lengths of RNA isolation techniques and their influence on mRNA degradation kinetics. (A) In vitro–transcribed mRNAs with defined poly(A) tail lengths were quantified by qPCR at the indicated site. RNA abundance was measured either by reverse transcription using oligo(dT) priming or after poly(A) selection with oligo(dT) beads followed by gene-specific priming. Values were normalized to input RNA (mean ± SEM, n = 3). (B) Quantification of poly(A) length with ePAT. The maximum-normalized gel intensity versus nucleotide length for PRO3 mRNA products indicates an average difference of 4 nts between the total and poly(A) RNA. **(C, D)** The impact of 5’–3’ degradation rates on the half-lives of the total and poly(A)^+^ mRNA in a model of mRNA decay. “Oligo(dT)” represents the sum of mRNAs with long and medium poly(A) tails, “LNA” includes mRNAs with long, medium and short poly(A) tails; “total” encompasses the entire mRNA population, including both poly(A)^+^ and poly(A)^-^ mRNA. The following parameter values were used γ_Exo_ = 0.025 min^-1^, *k_da1_* = 0.1, *k_da2_* = 0.5, *k_da3_* = 0.02 min^-1^. The fitted half-lives for the oligo(dT), LNA and total forms of the simulated unstable mRNA (γ_Xrn1_ = 0.5 min^-1^) were 1.1, 1.4 and 1.4 min, respectively (C); for the simulated stable mRNA (γ_Xrn1_ = 0.05 min^-1^), the corresponding values were 4.6, 12.3 and 12.3 min (D) (see also Fig. S4). **(E)** The steady-state fraction of short poly(A) tails in the total mRNA population as a function of mRNA half-life. The purple curve shows model predictions obtained by varying γ_Xrn1_ to modulate half-life. Gray arrows denote the increase in the fraction of short poly(A) tails after transcriptional shutoff for a duration equal to the half-life of the indicated mRNA. **(F)** Accumulation of short poly(A) tails during mRNA decay. The relative proportion of (A)_12_ poly(A) tails (mean ± SEM) was measured at steady state (0 min) and at the indicated time points following doxycycline-induced transcriptional shutoff (6 min for *DUT1*; 18 min for *PRO3*). Data represent P-values for the accumulation (difference between 0 min and the shutoff time points) were determined by Mann-Whitney U test (*N* = 3, *DUT1 P* = 0.7; *PRO3 P* < 0.05).

In yeast, the median poly(A) tail length is ∼26 nt (e.g., *PRO3*: 24 nt; *DUT1*: 31 nt) with the lengths of the individual molecules being broadly distributed^15^. Given this distribution, the ∼20 nt capture threshold identified above is expected to exclude a fraction of the mRNA population with short poly(A) tails. To test this, we measured poly(A) tail-length distributions using ePAT (Fig. S5A-E). Compared to poly(A)+ RNA, mean tail lengths in total RNA were ∼4 nt shorter for *PRO3* and ∼3 nt shorter for *DUT1* (Fig. 3B), supporting the model of preferential loss of transcripts with short poly(A) tails during isolation.

### Selective Loss of Short-Tailed Intermediates Leads to Underestimation of Stable mRNA Half-Lives

To determine how the capture threshold impacts half-life measurements, we built a mathematical model incorporating recent insights into mRNA processing (Equations (14) and (15)). The model assumes that decapping can occur during deadenylation without the need for complete tail removal. This is particularly relevant for stable mRNAs because they can be decapped allowing 5′–3′ degradation to begin even before deadenylation is complete^11^. The model also accounts for the nonlinear kinetics of deadenylation, which is initially slow, accelerates as tails shorten, and slows again when tails become very short^13^. Crucially, in this simulation, all mRNAs share the same deadenylation rate constants regardless of their overall stability. To compare the model with the experimental results, we defined three simulated RNA populations based on their tail-length composition: “oligo(dT)” (medium and long tails, at or above the (A)_20_ capture threshold), “LNA” (short, medium and long tails), and “total” (poly(A)^+^ and poly(A)^−^ RNAs). To assess how the degradation kinetics affects the estimated half-lives, we varied the rate of the 5’-to-3’ degradation since Xrn1 is the main mediator of differential mRNA stability^8^.

When the simulated degradation was fast, all three RNA populations yield similar half-lives (1.1 to 1.4 min) (Fig. 3C). In contrast, an mRNA parametrized with a 10-fold slower 5’-to-3’ decay showed a large discrepancy in half-life between the total or LNA population (12.3 min) and the oligo(dT) purified population (4.6 min) (Fig. 3D; Fig. S4A). The model thus reproduced the experimental findings and demonstrates that while deadenylation rates may be uniform, the underestimation of half-life caused by oligo(dT) isolation is largely restricted to stable mRNAs. Furthermore, the model confirms that LNA isolation effectively avoids these biases inherent to standard oligo(dT) beads.

While LNA-modified oligonucleotides effectively capture transcripts with short poly(A) tails, we reasoned that their performance might still diverge from total RNA measurements in genetic backgrounds where the relationship between deadenylation and decay is altered. To test this boundary, we examined mRNA half-lives in Δ*xrn1* cells, where the primary 5’–3’ exonuclease is absent and mRNA degradation is severely compromised. In this background, half-lives measured from total RNA were significantly longer than those from LNA-isolated mRNA, regardless of the transcript’s inherent stability in wild-type cells (Fig. S3).

This discrepancy can be explained by a model with an increased rate of terminal deadenylation (Fig. S4B), which may ensue due to the defects in mRNA dosage compensation in *Δxrn1* cells^24^. The accelerated terminal deadenylation leads to an accumulation of nearly tailless mRNAs that fall below the capture threshold of even LNA probes. To determine if this kinetic shift was dependent on translation, we measured, no-AUG mRNAs, in which translation was abolished by mutating all possible start codons. We again found a decreased half-life from the LNA-isolated mRNA, showing that translation does not alter this behavior in *Δxrn1* cells. These findings indicate that while LNA-based pull-down overcomes the primary biases of oligo(dT) beads in wild-type cells, its reliability as a proxy for total mRNA remains dependent on the presence of at least a minimal poly(A) tail, which may be lost in certain mutant contexts.

The above experiments show that the half-lives of stable mRNAs are more strongly affected by oligo(dT)-based isolation. A key implication of our model is that stable mRNAs progressively accumulate transcripts with shortened poly(A) tails, whereas accumulation is negligible for unstable mRNAs. Accordingly, the steady-state fraction of mRNAs with short poly(A) tails is expected to increase with mRNA half-life and to rise further during decay following transcriptional shutoff (Fig. 3E). These short-tailed transcripts are precisely the population expected to be excluded during standard poly(A)+ mRNA isolation.

While previous studies have reported that specific mRNAs accumulate short poly(A) tails of ∼10–12 nt^25^, our model predicts that this is a general phenomenon. To test this, we developed a modified ePAT-based assay (ePAT–qPCR, Fig. S5F) to selectively detect short poly(A) tails (∼A₁₂) in defined UTR isoforms. We then compared the proportion of short poly(A)-tailed transcripts (Equation (4)) for *DUT1* and *PRO3* following transcriptional shutoff. At 6 and 18 min after doxycycline addition, the overall mRNA levels of both transcripts declined similarly. However, the fraction of short poly(A)-tailed mRNAs increased significantly only for the relatively stable *PRO3* mRNA (Fig. 3F), in agreement with the model predictions.

### Systematic Overestimation of Translation Efficiency in Poly(A)-Selected Datasets

The above results confirmed that the estimated half-lives of mRNAs with high codon optimality are particularly sensitive to the type of RNA isolation. Following this validation, alternative hypotheses remain to explain the weaker correlation between CTCs and tRNA copy number in total RNA compared to poly(A)+ RNA.

First, mRNA abundances in the Riba RiboZero dataset were obtained from a different study^17,26^, which may have introduced experimental noise or reflected specific conditions where stability and translation are regulated independently. Alternatively, the discrepancy may arise from how half-lives affect steady-state mRNA levels. To quantify this, we applied the kinetic model from Fig. 3 to equilibrium conditions. This steady-state solution (Fig. 4A) shows that stable mRNAs accumulate a larger fraction of deadenylated intermediates that are missed by poly(A)+ selection. Since TE is calculated as the ratio of protein production rate to mRNA abundance, this underestimation of mRNA levels causes an artificial overestimation of TE for stable mRNAs.

**Figure 4.**
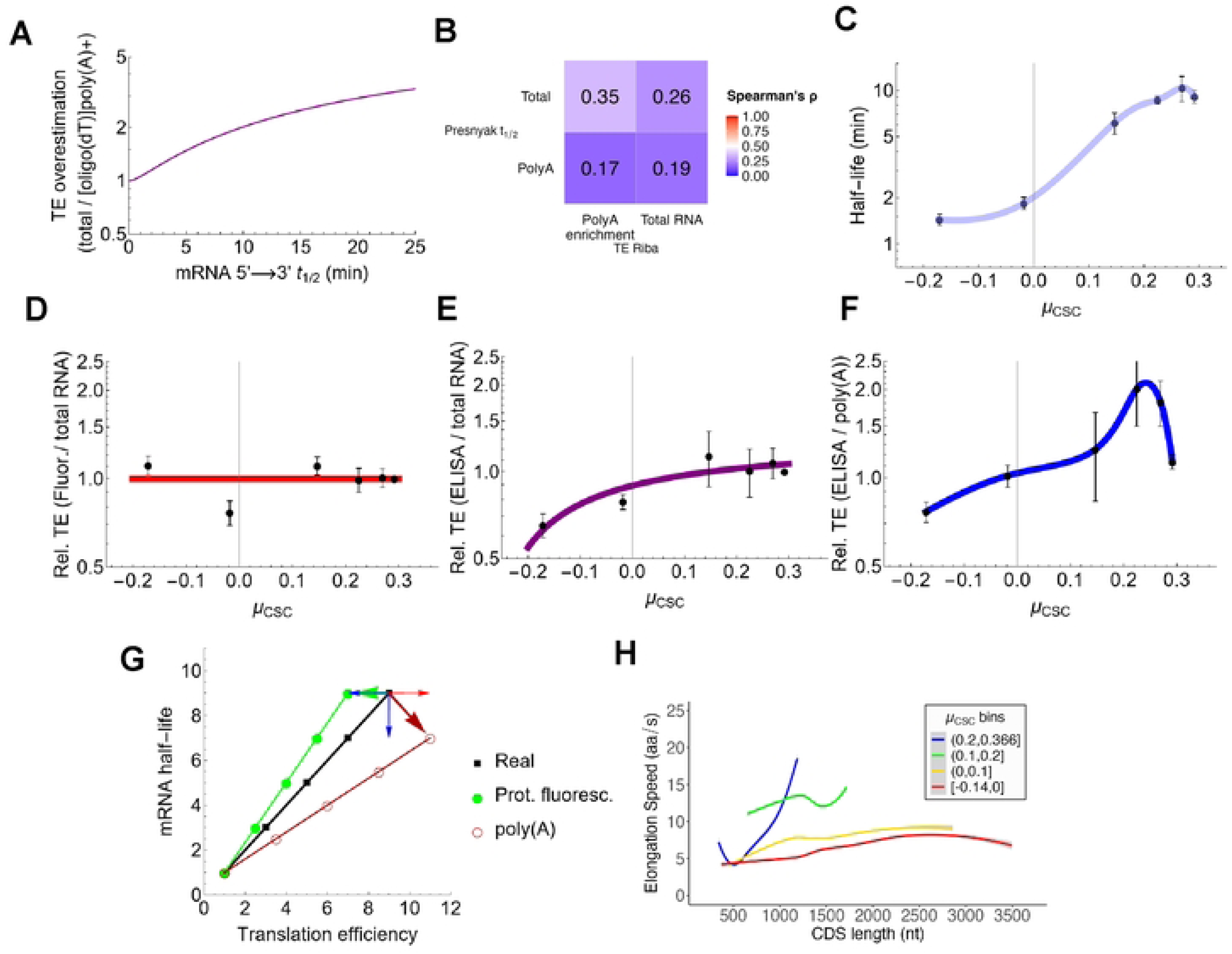
Impact of mRNA isolation and protein quantification methods on translation efficiency (TE) estimates. **(A)** Mathematical model showing the predicted overestimation of TE when using poly(A)+ selection. The panel shows the steady-state solution of the model presented in Fig. 3C, D. The steady-state mRNA ratio (total/oligo(dT)) is plotted as a function of mRNA half-life (determined by t_1/2_ = Ln(2) / γ_Xrn1_ ). Model parameters: γ_Exo_ = 0.025 min^-1^, *k_da1_* = 0.1, *k_da2_* = 0.5, and *k_da3_* = 0.02 min^-1^. **(B)** Correlation matrix (r_s_) between the Presnyak mRNA half-life datasets with eliminated outliers and the Riba TE datasets with poly(A) and total mRNA isolation. **(C)** Measured half-lives of GFP–RFP reporter mRNAs with varied codon optimality (µ_CSC_). Error bars: SEM (n = 3 replicates). **(D-F)** Relative TE calculated for recoded GFP-RFP mRNAs comparing: fluorescence and total RNA (D), ELISA and total RNA (E), and ELISA and oligo(dT)-isolated mRNA (F). Error bars denote SEM (*n* = 3). **(G)** Vector scheme illustrating how technical biases in measurement (fluorescence maturation and mRNA capture efficiency) shift perceived TE and half-life away from real biological values. **(H)** Elongation speed of the yeast transcriptome calculated based on Riba (RiboZero) dataset (1099 mRNAs). Curves represent moving averages (n=50) for mRNAs binned according to µ_CSC_ ranges.

We examined the relationship between TE and half-lives as a function of RNA isolation method across the transcriptome, using a filtered dataset for mRNA half-lives (Fig. S2). Specifically, half-lives measured from total RNA correlated more strongly with TEs derived from poly(A)-selected RNA than from total RNA (Fig. 4B). Conversely, poly(A)+-based half-lives correlated better with TEs calculated from total RNA than from poly(A)-selected RNA. This cross-dependence is particularly striking given that the TE values originated from independent datasets generated in different studies and therefore subject to distinct experimental noise. Thus, combining different datasets does not necessarily amplify noise to mask true relations.

To experimentally confirm that cross-dependence of the parameters with the isolation methods, we constructed a series of mRNAs encoding a tandem GFP–RFP fusion protein with varying degrees of codon optimization. mRNA half-lives were measured using multiplexed gene control methods. As expected, half-lives increased from 1.4 to 10.4 minutes with rising codon optimality (Fig. 4C). Interestingly, the construct with the highest proportion of optimal codons showed a slight decline in half-life, suggesting that stabilization does not further improve once a threshold of codon optimality is exceeded, a phenomenon suggested in the context of translation^27^.

To estimate TE based on steady-state protein to mRNA ratios, we measured fluorescence of GFP and RFP. Additionally, we measured the decay rates of GFP–RFP by monitoring fluorescence after shutting off gene expression using flow cytometry. The proteins were stable and exhibited indistinguishable half-lives, irrespective of codon optimality (Fig. S6A) This uniform protein stability justifies our use of steady-state protein-to-mRNA ratios as a direct proxy for TE (Equation (12)). When mRNA was quantified from total RNA, TE appeared nearly constant and largely unaffected by codon optimality (Fig. 4D). Given this unexpected lack of effect, we repeated the analysis using RNA isolated with oligo(dT) beads. Under these conditions, TE showed greater variability: mRNAs with the four highest µ_CSC_ values followed the same trend as half-lives, yet overall, there was no clear separation between mRNAs with negative versus positive codon optimality, resulting in a fluctuating pattern (Fig. S6B).

We reasoned that this discrepancy arose because high translation rates can exceed the folding capacity of GFP and RFP. Since codon optimality does not affect protein stability (Fig. S6A), the observed discrepancy in fluorescence levels likely arises from differences in fluorophore maturation rather than degradation. To confirm this, we cultured cells at a lower temperature (20°C) to alleviate folding bottlenecks caused by rapid translation. We reasoned that fluorophore folding would be less affected by reduced temperature than translation because it is a non-catalyzed process. In contrast, enzymatically-catalyzed reactions like translation would be expected to decline more steeply as temperature decreases^8^. Indeed, lowering the temperature led to a twofold increase in fluorescence for constructs with positive compared to negative µ_CSC_. (Fig. S6C). In addition, we used ELISA targeting GFP as a fluorescence-independent measure of protein abundance. Fluorescence quantifications yielded results highly consistent with ELISA at 20°C (Fig. S6C), indicating that decreasing the translation rate alleviated the deficit in fluorophore maturation.

To circumvent this limitation of fluorophore formation at standard temperature (30°C), we directly quantified protein abundance by ELISA. The ratio of fluorescence to ELISA signal decreased sharply as µ_CSC_ became positive (Fig. S6D), confirming that faster elongation rates impair fluorophore maturation. Notably, the ELISA-based TE increased monotonically with codon optimality, declining only slightly in the construct with the highest µ_CSC_ (Fig. 4E). The overall dependence resembled a hyperbolic, Michaelis–Menten–like function, revealing a relationship between codon optimality and TE that had been masked by fluorescence-based measurements. Notably, the range of TE values was less than twofold—much narrower than the range observed for half-lives. However, when ELISA was combined with RNA isolated via oligo(dT) beads, the TE range expanded (Fig. 4F), and the dependence of TE on codon optimality more closely paralleled that of half-lives (Fig. 4B); however, this increased TE again confirms the bias due to the inefficient capture of poly(A) mRNAs (Fig. 4A).

Together, these results demonstrate that the apparent dependence of TE on codon optimality is reduced when total RNA is used instead of poly(A)+ RNA. This inverse behavior relative to mRNA half-lives cannot be explained by dataset-specific noise (Fig. 4B) or distinct conditions or regulatory mechanisms. Rather, it reflects an overestimation of TE caused by the inefficient capture of mRNAs during poly(A)+ selection, leading to an artificially small denominator in TE calculation (Fig. 4A, G).

### Codon optimality enhances ribosome loading and accelerates elongation speed

By combining TE datasets with polysome profiling measurements of mean ribosome occupancy (MNR)^16^, we inferred ribosome flux (elongation speed, Equation (13)). The mean elongation speed increased two- to threefold across codon optimality bins (Fig. 4H) and, at high codon optimality (μ_CSC_ > 0.2), showed a strong dependence on CDS length. Notably, the effect of codon optimality on elongation appears to plateau near the average CDS length. This phenomenon coincides with shifts in codon composition close to the average CDS length. There are very few genes above 1500 nt that possess highly optimal codons and, consequently, rapid elongation (Fig. 4H). Thus, the slower translation and instability of many long mRNAs appear to reflect an enrichment in non-optimal codons with increasing CDS length, rather than a distinct molecular mechanism, consistent with a trend observed across multiple model organisms that frequency of optimal codons decreases with the length of the encoded protein^28^.

To elucidate how TE and elongation speed responds to codon optimality, we analyzed ribosome distributions. We performed polysome profiling of the GFP–RFP expressing strains and quantified mRNAs associated with zero, one, two, or more ribosomes (Fig. 5A). The proportion of ribosome-bound mRNAs (fractional occupancy, Equation (1)) increased sharply between μ_CSC_ values of –0.2 and 0.1, after which it plateaued at ∼95% (Fig. 5B). This pattern closely mirrored the transcriptome-wide trend (Fig. 5C) as our analysis of previously published data indicates. This suggests that codon optimality in this range enhances ribosome recruitment to mRNAs.

**Figure 5.**
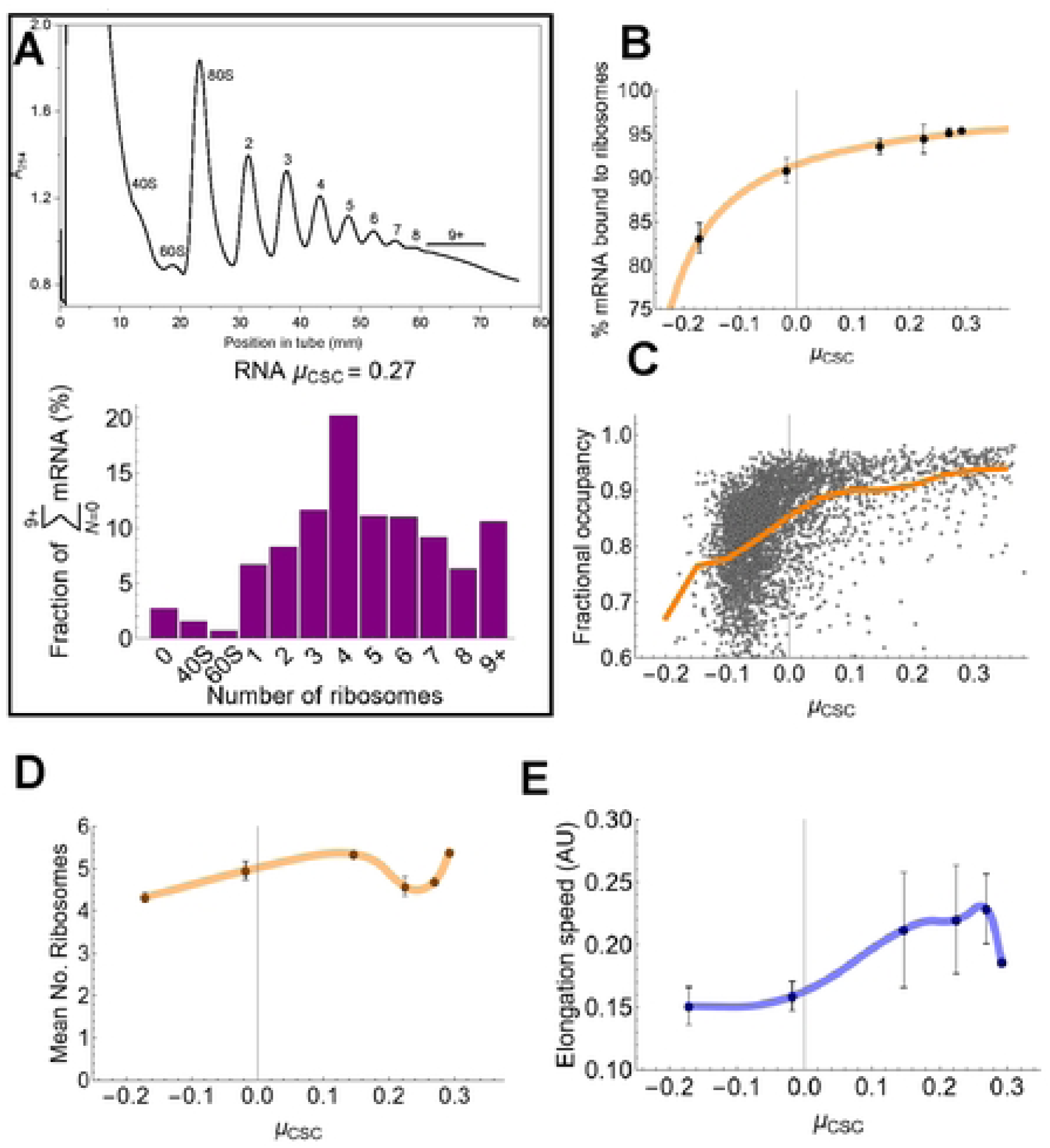
Translational capacity of mRNAs as a function of codon optimality. **(A)** Polysome profiling was performed to separate mRNA fractions based on ribosome occupancy (top panel). mRNA abundance in each fraction is expressed as a fraction of the total mRNA across all detected ribosomal states (0,40S,60S, and 1 to 9+ ribosomes; shown here for the G-Rp027 mRNA with µ_CSC_ = 0.27. **(B).** Ribosome-bound mRNA (in %, corresponding to 100× fractional occupancy) of GFP-RFP CDSs recoded with the indicated µ_CSC_ values. Error bars represent SEM (*n* = 3 replicates). **(C)** Fractional occupancy of the yeast transcriptome. The orange line connects mean values within the indicated µ_CSC_ ranges. **(D)** Mean number of ribosomes associated with the mRNAs shown in Fig. 3. **(E)** Relative elongation speed of the mRNAs shown in (D), calculated as v = ρ · TE, where ρ = MNR / CDS length.

We next examined the polysome propensity, defined as the ratio of mRNAs bound by three or more ribosomes relative to those bound by a single ribosome. This metric displayed a non-monotonic dependence: it increased initially, dipped around μ_CSC_ ≈ 0.2, and then partially recovered (Fig. S6E)—a trend also evident at the transcriptome level (Fig. S6F). These results indicate that while moderate codon optimality promotes polysome formation, higher values may impose kinetic constraints that reduce ribosome loading efficiency^29^.

Together, these parameters shape the mean number of ribosomes (MNR, Equation (2)) associated with mRNAs, which likewise rose initially but dropped near μ_CSC_ ≈ 0.2 (Fig. 5D). From MNR and translation efficiency values, we inferred elongation speed, which increased with μ_CSC_ (Fig. 5E) but declined at the highest values, mirroring the behavior of mRNA half-life.

### Assessment of translation capacity during mRNA aging

Having characterized how codon optimality dictates ribosome recruitment and speed, we next asked whether this translational capacity is maintained throughout a transcript’s lifecycle. This is a critical question because the apparent overestimation of translation efficiency with poly(A)+ mRNA might be mitigated if transcripts with short poly(A) tails are translated less efficiently or not at all (Fig. 6A).

**Figure 6.**
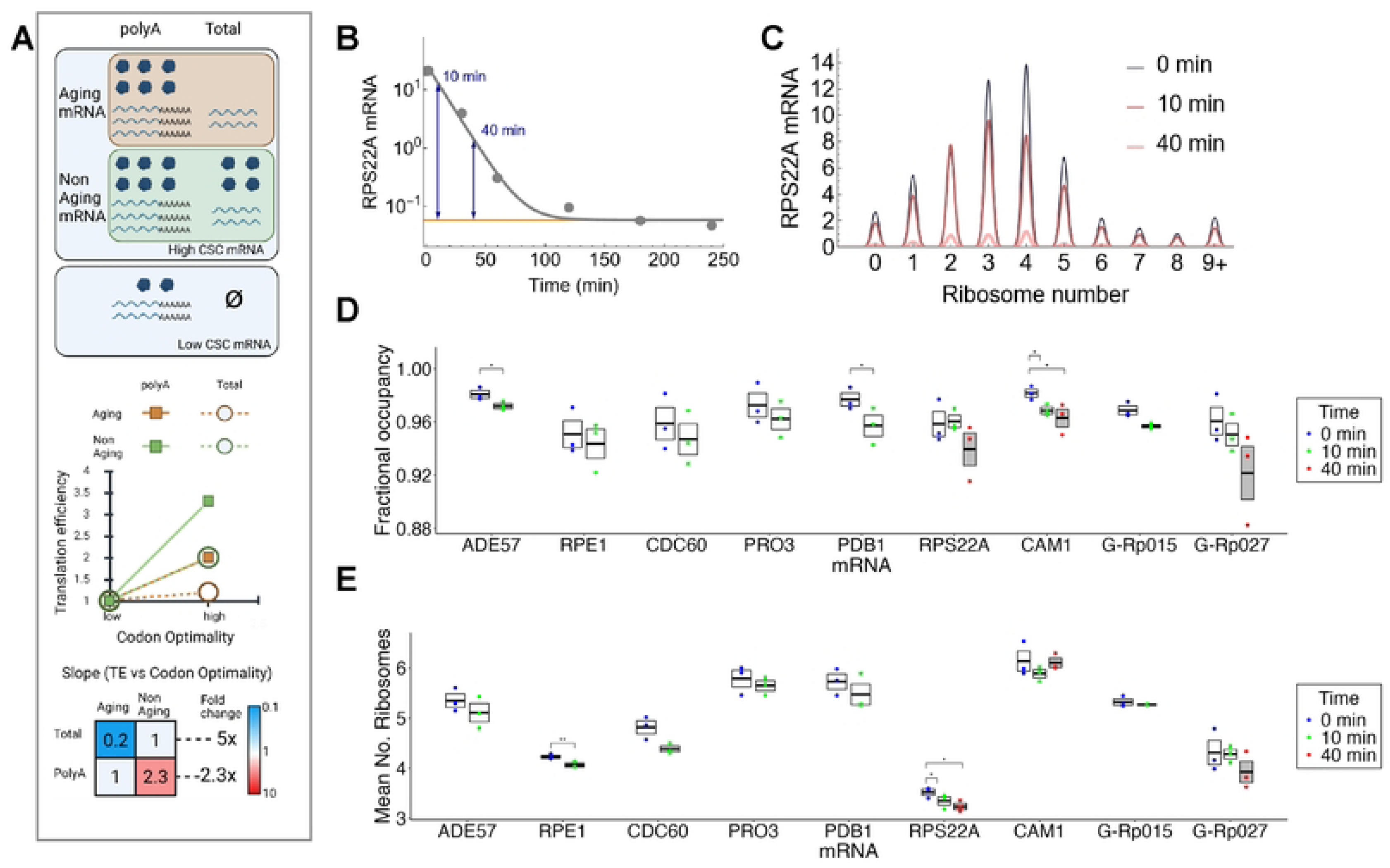
**Relationship between mRNA aging and translational capacity**. **(A)** Scheme showing how mRNA aging influences translation efficiency measurements and how different isolation methods affect results. Top: mRNA aging model: poly(A)-tailed mRNAs are translated, while those lacking tails are not, impacting high-CSC but not low-CSC mRNAs. Middle: TE vs µ_CSC_ with or without poly(A) selection and mRNA aging. Bottom: Slopes of TE vs µ_CSC_ under different aging conditions and isolation methods. **(B)** Estimation of mRNA half-life and residual baseline. RPS22A levels (gray dots) were fitted with an exponential decay model including a nonzero baseline (*b*) (gray curve). The orange line marks the baseline level (*b*) determined from a long, 4-hour decay experiment. Blue arrows indicate the ratios of RNA abundance to the baseline at 10 and 40 minutes. At these time points, RPS22A was 897- and 32-fold above baseline, respectively. **(C)** Abundance of RPS22A mRNA across polysome fractions during transcriptional shut-off. The peaks of the bell-shaped curves represent mRNA levels at each ribosome number at 0, 10, and 40 min after doxycycline addition. **(D, E)** Fractional occupancy (D) and mean number of ribosomes (E) associated with mRNAs 0, 10 and 40 min after addition of doxycycline. *N* =3 replicates. * and ** show significance of differences at p < 0.05 and p < 0.01, respectively.

To test this possibility, we measured fractional ribosome occupancy and the MNR after shutting off gene expression with doxycycline, thereby preventing transcript replenishment and allowing the isolation of aging transcripts. Samples were collected before and 10 and 40 minutes after shutting off gene expression, followed by polysome profiling. We focused our analysis on a set of endogenous mRNAs spanning a wide range of total-to-poly(A)+ half-life ratios (Fig. 2B), as these transcripts represent varying degrees of potential poly(A)+ selection bias. Across these targets, as well as two recoded GFP-RFP constructs, we generally observed only a slight decline in both occupancy and MNR in some of the aging transcripts.

As an illustration, RPS22A abundance decreased across all fractions after 10 and 40 minutes (Fig. 6B, C); however, the resulting change in translation parameters was small, typically less than 10%. Thus, while a stable mRNA may suffer from a 1.5–3-fold underestimation of steady-state abundance in poly(A)+ datasets due to its half-life ratio, we observed only a marginal ∼1.1-fold decline in translation capacity for those same aged transcripts (Fig. 6D, E). Since the technical error in the denominator is significantly larger than the physiological shift in the numerator, the distortion in TE estimates is not meaningfully mitigated by mRNA aging.

### Contribution of codon optimality to translational efficiency and half-lives as a function of CDS length

Codon optimality parameters such as the tRNA adaptation index (tAI) correlated more strongly with half-life (ρ = 0.61) than with TE (ρ = 0.49) (Fig. S1E). However, correlation strength does not necessarily reflect control strength. Control strength measures how steeply an output variable changes as an input variable varies, whereas correlation conflates slope and measurement noise. The lower TE correlation could therefore reflect higher noise, since fewer datasets contributed to the unified TE values than to the half-life dataset (3 versus 8).

To compare the impact of codon optimality on half-life and TE, we calculated control coefficients using Theil–Sen robust regression. This approach is less sensitive to outliers than ordinary least squares^30^. We calculated the slopes between the tAI, which reflects tRNA gene copy numbers and codon–anticodon interaction strength^31^, and the TE or half-life normalized by their respective medians (Fig. 7A). Steeper slopes indicate greater control by codon optimality. While half-life control coefficients remained relatively constant across length scales, TE coefficients varied significantly: for short CDSs (<400 nt), the control of stability and translation were similar, but for average-length mRNAs, TE control began to exceed that of half-life (Fig. S7A).

**Figure 7.**
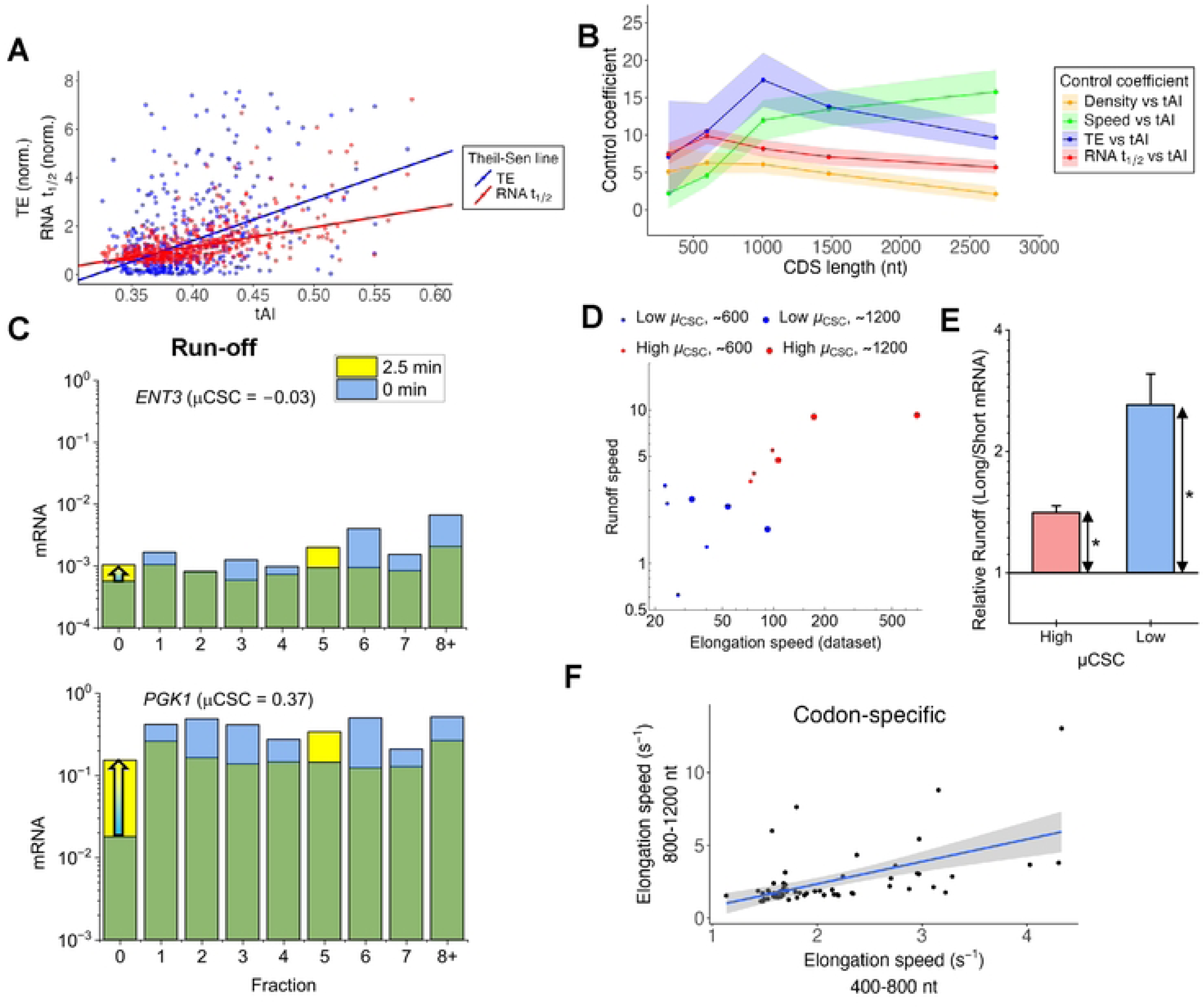
Transcriptome wide dependence of half-lives, elongation speed and translation efficiency on codon optimality. All analyses were performed using a unified TE dataset extrapolated to total mRNA. **(A)** The dependence of *t_1/2_* and TE on tAI of the mRNAs with CDS between 800 and 1200 nts. **(B)** Control coefficients for mRNA stability, translation efficiency, elongation speed and ribosome density as a function of CDS length. Shaded areas indicate 95% confidence intervals. **(C)** Ribosome run-off for ENT3 and PGK1 (CDS: 1227 and 1251 nt). Normalized mRNA abundances across polysome fractions at 0 and 2.5 min after glucose depletion. Arrows indicate a 1.84-fold increase in the free fraction (0) for *ENT3* versus 8.58-fold for *PGK1*, reflecting faster elongation in the more optimal transcript. **(D)** Comparison of elongation speed estimated from TE datasets and ribosome runoff. mRNAs with low variance in TE and having shorter (∼600 nt, small circles) and longer (∼1200 nt, large circles) CDSs with low (blue) and high (red) codon optimality were selected. **(E)** Relative runoff was calculated as the ratio of runoff speeds between the full-length (1467 nt) GFP-RFP constructs and corresponding 735-nt fragments. Ratios were significantly greater than 1 for both high and low codon optimality (Ratio t-test, p=0.013 for high µ_CSC_, p=0.047 for low µ_CSC_). Error bars denote SEM (N=3). **(F)** Comparison of codon-specific elongation speeds (CS) calculated for mRNAs with different CDS length ranges (400-800 and 800-1200 nt; 422 and 471 genes respectively) from the unified TE dataset normalized to extrapolated total mRNA. Linear regression yielded an adjusted R² = 0.42 and Spearman’s ρ = 0.80. The shaded area denotes the 95% confidence interval of the regression line.

While these results are based on total RNA (Riba RiboZero dataset), they are limited in coverage to approximately one thousand mRNAs. To extend our analysis to more mRNAs, we used the unified TE dataset, which covers more than half of the transcriptome. Because this dataset is poly(A)-based rather than total RNA-based, we converted the poly(A) TE values to predicted total TE values using regression (Equations (8), (9)). Specifically, we derived a regression function between the poly(A)-to-total mRNA ratio and the corresponding half-life ratio from the Presnyak dataset (Fig. S7B). This approach yielded control coefficient dependencies similar to those obtained from RiboZero data (Fig. S7C): the TE control coefficient was approximately threefold higher than that for half-life when using poly(A)⁺ measurements, but decreased to roughly twofold when using the regression-based extrapolated total mRNA values (Fig. S8).

The resulting control coefficients displayed trends consistent with those observed in the smaller dataset. For shorter mRNAs (<800 nt), control coefficients for TE and half-life were indistinguishable (P = 0.86 and 0.57; Fig. 7B). However, for longer mRNAs—particularly those around 1000 nt, which is closer to the average transcriptome-wide CDS length (∼1500 nt) — the TE control coefficient was approximately twice that of half-life (P = 10^-4^, permutation test). This indicates that in longer transcripts, codon optimality exerts stronger control over translation than decay.

### Contribution of the control by elongation and ribosome density

This increase in translational control is paralleled by a similar increase in the control of elongation speed, whereas the contribution of ribosome density remains low and constant across CDS lengths (Fig. 7B). To independently verify these elongation speed estimates, we performed ribosome run-off assays. Upon abrupt glucose depletion, translation initiation is inhibited, allowing elongating ribosomes to run off from mRNAs ^32^. The extent of run-off—quantified as the fold change in mRNA abundance in the free fraction after 2.5 min—provides an estimate of elongation speed (Equation (3)). For example, *PGK1* mRNA (1251 nts) exhibited a 5.5-fold faster run-off than the *ENT3* mRNA, which has a similar length (1227 nts) but lower codon optimality (Fig. 7C). A similar, 7.6-fold faster elongation speed was estimated for *PGK1* based on TE and ribosome density. Systematically comparing mRNAs across the relevant CDS length ranges with consistent estimates across both methods revealed a strong correlation (Fig. 7D, Spearman’s ρ = 0.71, P = 0.005).

In addition, we compared the run-off speed of synthetic mRNAs of different lengths, namely full-length GFP–RFP constructs and corresponding half-length fragments. Run-off speed was significantly higher for the longer mRNAs, independent of codon optimality (Fig. 7E), indicating a length-dependent effect on elongation dynamics.

Beyond elongation rate, codon optimality has been recently suggested to positively influence the initiation rate^33,34^. which can be qualitatively assessed by measuring ribosome density. Our data shows that ribosome density is positively controlled by codon optimality, though less strongly than elongation (Fig. 7B). However, whether the effect of codon optimality on initiation is more or less significant than its effect on elongation remains dependent on the mechanism, specifically regarding whether translational re-initiation occurs through mRNA circularization^35^.

If ribosomes are fully recycled from the 3’ to the 5’ end, the ribosome density (ribosomes per transcript) remains independent of elongation speed and codon optimality because the total number of ribosomes in the circuit is conserved. In contrast, if translation is initiation-limited, an increase in elongation speed (driven by higher codon optimality) causes ribosomes to move through the CDS more quickly, which would naturally tend to decrease ribosome density. Consequently, in this second scenario, the inferred feedback effect of codon optimality on initiation must be much stronger to overcome the density-reducing effect of faster elongation and produce the net increase in density observed in our data (Fig. 7B). Determining feedback strength independently of the model typically requires specialized methods, such as feedback opening^36^.

Finally, we focused on the range of mRNAs exhibiting the highest TE control coefficients to derive precise codon-specific elongation rates via Bayesian analysis. Estimated speeds ranged from ∼1 to 15 codons per second (median ∼2 codons/sec, Fig. S7D). Comparing codon-specific elongation rates between shorter and longer CDS bins yielded a linear relationship of *l* = 1.5*s* – 0.66, where *s* represents the 400–800 nt bin and *l* the 800–1200 nt bin (Fig. 7F). This trend suggests that increased elongation rates at a subset of optimal codons in longer CDSs may contribute to the stronger elongation control observed in these transcripts.

## DISCUSSION

Post-transcriptional regulation often involves the tight coupling of mRNA translation and degradation. In many regulatory contexts—such as microRNA-mediated repression—a central debate persists regarding whether translation inhibition or accelerated mRNA decay is the primary driver of the observed gene downregulation^37–39^, with some works suggesting a predominant effect on mRNA half-lives.

While the coupling of these processes through codon optimality is now well-recognized, the relative contribution of translation efficiency versus mRNA stability to overall protein output has remained less clear. Our results demonstrate that measurement strategies play a decisive role in this determination. Specifically, we show that the choice of protein detection method and mRNA isolation technique can systematically distort the perceived relationship between codon optimality, half-life, and translation.

Two main methods are commonly used for quantification in proteomics: fluorescent reporter assays and mass spectrometry^40^. We were surprised to find that fluorescence-based measurements of translation efficiency did not detect changes in codon optimality. In contrast, switching to direct quantification by ELISA revealed such changes. This discrepancy likely reflects the fact that codon optimality can influence folding speed, and proteins may adopt alternative conformations^41–43^. GFP and other fluorescent proteins undergo several maturation steps before becoming fluorescent, including folding and oxidation–reduction reactions. If elongation speed outpaces folding and fluorophore maturation, then fluorescence will underestimate protein output. Consistently, we observed improved translation efficiency measured by fluorescence when cells were grown at 20 °C, a condition that slows elongation. Similarly, bacterial studies have shown markedly improved folding and fluorescence at reduced temperatures, which compensate for the inherently faster elongation rates in bacteria compared to eukaryotes^44,45^. Other biophysical factors may also contribute^46^. Evolutionary constraints likely underpin this sensitivity, as many fluorescent and luminescent proteins originate from organisms adapted to the cooler environments of deep-sea water, where maturation kinetics are optimized for lower temperatures.

While fluorescence remains a valid tool for comparing the same protein across constant conditions, experiments altering synonymous codon usage must be interpreted with caution. Because ELISA and mass spectrometry directly quantify protein abundance, they are not confounded by folding efficiency and thus provide a more accurate conclusion regarding the codon-to-translation relationship.

While fluorescent readouts can underestimate translation efficiency—an effect most pronounced at high codon optimality—the isolation of mRNA with oligo(dT) beads has the opposite effect, artificially amplifying TE. Oligo(dT)-based isolation became common with the advent of RNA-seq–based quantification. Earlier methods such as qPCR and DNA microarrays directly quantified mRNAs using sequence-specific oligonucleotides, whereas RNA-seq sequences all RNAs in a sample, the majority of which are rRNAs. Although rRNA can be depleted, depletion is species-specific, variably efficient, and more costly^47^; therefore, oligo(dT) isolation remains more widely used. Each method has particular strengths for capturing different parts of the transcriptome^48^, but studies that rely on poly(A)-based processes are especially vulnerable to biases introduced by poly(A) selection^49^. Methods have been developed to mitigate the biases by improving the representation of mRNAs upon poly(A) selection^50^. We found that bias in half-life measurements arises only from mRNA isolation with oligo(dT) beads, not from oligo(dT) priming during qPCR. While bead-based isolation requires a minimum tail length of ∼20 nt, oligo(dT) priming effectively captures transcripts with even shorter or absent poly(A) tails. This robustness in priming likely arises due to internal priming at A-rich tracts or template-switching^51–54^.

Because steady-state distributions of poly(A) tail lengths are shaped by differential half-lives, stable mRNAs are underrepresented in oligo(dT) capture. As a result, translation efficiency estimates become inflated. Crucially, this creates a zero-sum trade-off in the perceived control of gene expression: any bias that artificially improves the correlation between codon optimality and TE simultaneously deteriorates its correlation with mRNA stability, and vice versa.

It is likely that this bias represents a conserved phenomenon, as mRNAs across most organisms—including humans—harbor a substantial proportion of short poly(A) tails^15^. In particular, cell-cycle regulatory genes undergo dramatic deadenylation in M-phase, which serves as a primary mechanism for their translational repression^55^. Beyond tail-length dynamics, recent studies in specialized cell types, such as macrophages, have identified unconventional codon usage as a key mediator of translational dynamics of cell cycle related mRNAs^56,57^. Future studies will be required to determine to what extent these effects are driven by tRNA abundance^58^. Ultimately, these observations suggest that poly(A) tail dynamics are expected to significantly influence human translatome estimation, particularly for transcripts with highly dynamic tail-length distributions such as those regulated during the cell cycle.

The biases introduced by oligo(dT) capture could be in principle mitigated by mRNA aging. However, our results indicate that mRNA aging affects the translation capacity of few mRNAs and, even for those, the effect is much smaller than the bias introduced by the half-life shift (Fig. 6D, E). This is reminiscent of prior studies in mammalian cells that did not detect mRNA aging with synthetic constructs, whereas some endogenous mRNAs were found to display aging and aging-dependent reductions in translation^59,60^. This suggests that the phenomenon may be mRNA- and condition-specific. The detectable nature of this process, even though minor in scale, indicates that it may be non-negligible for specific mRNAs and should be considered in future studies.

The RNA isolation method also affects the estimated control coefficients. While the TE control coefficient was approximately threefold higher than that for half-life when using poly(A)⁺ measurements, this decreased to roughly twofold when using regression-based extrapolated total mRNA values (Fig. S8). This demonstrates that while translational control remains dominant, but its estimated magnitude is reduced after accounting for poly(A)-selection bias.

We found a clear CDS-length threshold for codon-optimality–dependent control of TE, but not for mRNA half-life, over the examined length range. The TE control coefficient increased with CDS length up to ∼1000 nt and reached approximately twice the value of the half-life control coefficient at this length. By contrast, the half-life control coefficient remained comparatively constant across CDS lengths. However, this does not rule out half-life thresholds for mRNAs with very short CDSs. In fact, our earlier study with experimentally recoded genes suggested a threshold for codon optimality effects on mRNA half-life between 200 and 400 nt^16^. Whereas half-lives of short mRNAs are not or are minimally affected by codon recoding, the response emerges above this threshold. In the present datasets, the lowest bin overlaps with the upper limit of this threshold, and because very few mRNAs fall below this range, this threshold could not be resolved. Thus, the threshold for half-life control likely lies at a shorter CDS length than that for translational control. The observed length dependence of TE control may reflect CDS length–dependent re-initiation of terminating ribosomes^61^, or other translation-specific processes that do not affect decay. Because measurement strategies can substantially shift the apparent balance between these processes, integrated measurement approaches are essential to accurately dissect their relative contributions.

## ACKNOWLEDGEMENTS

We thank Gowtham Thambra, Ema-Andreea Palastea for the for experimental help, Sylvia Voegeli for plasmids, Phillippe Demougin (Genomics Facility, University of Basel / BSSE) for the TruSeq mRNA isolation.

## AUTHOR CONTRIBUTIONS

A. B., S.R., S.M. and M.B. designed the experiments and data analysis. S.R., C.E.D. M.B, S.W. and C.P performed the experiments. A. B. wrote the manuscript. S.M., A.B., S.R. and V.J. analyzed the data.

## Data Availability

All relevant data are within the paper and its Supporting information files.

## SUPPLEMENTAL INFORMATION

Figures S1-S8

Table S1 Translation efficiency (TE) datasets used for the analysis and dataset unification.

Data S1 Data related to computational analysis, underlying Figs. 1 and S1, including P-values

Data S2 Data related to poly(A)-dependence of decay kinetics, underlying Figs. 2 to 3 and S2-S5

Data S3 Data related to translation efficiency and elongation, underlying Figs. 4 to 7 and S6-S8

Data S4 Unified TE dataset and related parameters

Data S5 Tet-controlled strains

Data S6 ePAT-qPCR primers Data S7 -RT control

## METHODS

### Cell culturing conditions

For measurements at 30°C, *Saccharomyces cerevisiae* BY4743 isogenic cells were inoculated and grown overnight at 30°C.

For the determination of TE, they were then refreshed to an OD600 of 0.05 and subsequently grown at 30°C until the mid-log phase (OD600 ∼ 0.5). This ensured that three cell divisions were completed to reduce carryover of protein produced in the stationary phase.

For the mRNA decay experiment, yeast cells expressing a gene of interest (Data S5) under the control of TET-off system were used. Overnight cultures were refreshed to an OD600 of 0.15 and subsequently grown at 30°C until the mid-log phase (OD600 ∼ 0.5). The expression of genes under TET-promoters was halted by addition of doxycycline (Sigma-Aldrich #D9891) to a final concentration of 10 μg/ml. Samples were taken at specific time points to study the decay profile of the relevant mRNAs as previously described^16^.

For the determination of TE at 20°C, cells were inoculated and grown 30°C for 6-8 hours, following which the cultures were grown overnight at 20°C. They were then refreshed to an OD600 of 0.05 and subsequently grown at 20°C until the mid-log phase (OD600 ∼ 0.5). This typically takes ∼20 hours.

Extraction of total RNA

5mL or 50mL (depending on the amount of RNA needed) of the yeast culture in the mid-log phase was collected in chilled methanol (final concentration 50% v/v). The resulting mixture was centrifuged at 2500g, 4°C for 10min and then the supernatant was removed. Total RNA was extracted using RNeasy Mini or Midi Kit (Qiagen #74106, #75144) according to kit protocol with the addition of 1% β-mercaptoethanol in buffer RLT, including an on-column DNase digestion step,. Lysis was carried out using appropriate MPBio Lysing Matrix Y tubes (MPBio #116960500 or #116975050) in a FastPrep-24™ 5G bead beating grinder and lysis system (MPBio #116005500).

Isolation of poly(A)+ RNA

Poly(A)+ RNA was isolated with the Poly(A)Ttract mRNA isolation Systems III or IV (Promega , abbreviated as Method P) or TruSeq Stranded mRNA Library Prep (96) [20020595] (Illumina, abbreviated as Method TS). When the oligo(dT) was replaced by LNA-oligo(T), the PolyATtract protocol was modified: the washing and elution was made more stringent, by adding sarkosyl to washing steps and the elution was performed with distilled water preheated to 65°C instead of room temperature.

In brief, 500 μl of the total RNA (250 - 500 ng/µl), obtained with the RNeasy Midi , was warmed to 65°C for 10 minutes, then mixed with 0.3 μM of biotinylated- LNA-oligo(T) Probe and 13μl of 20X SSC solution (Promega), and incubated at room temperature for 10 minutes. This solution was then transferred to the tube containing washed Streptavidin Paramagnetic Particles, and incubated for 10 minutes. The beads with the annealed mRNA were washed twice with 0.1x SSC solution containing 0.1% N-Lauroyl Sarcosine, and twice with 0.1x SSC solution. The mRNA was eluted by re-suspending the washed pellet in 100 μl of the RNase-Free water pre-warmed to 65°C, followed by an incubation at 65°C for 10 minutes and a rapid transfer to ice for 5 minutes. Finally, the supernatant containing the mRNA was separated from the recaptured particles, and was supplemented with RNasin® Plus RNase inhibitor (Promega).

### Polysome profiling during steady state and mRNA aging conditions

For polysome profiling during steady state conditions, yeast cells were collected in the mid-log phase as previously described^16^.

For polysome profiling during mRNA aging conditions, yeast cells expressing a gene of interest under the control of TET-off system were used. When the cultures were in the mid-log phase, gene expression was halted by addition of doxycycline (Sigma-Aldrich #D9891) to a final concentration of 10 μg/ml, and samples were collected at 0 minutes (prior to doxycycline addition), 10 minutes and 40 minutes, following which polysome profiling was carried out.

For the polysome profiling, yeast samples were collected by adding 1 volume of culture to 2 volumes of ice-cold media with a final concentration of 200 µg/mL cycloheximide (CHX) to arrest translation. Cells were pelleted by centrifugation and all traces of supernatant were removed. Cell pellets were immediately lysed in PLB (20 mM Tris-HCl pH 8.0 (Invitrogen #AM9855G), 10 mM MgCl_2_ (Invitrogen #AM9530G), 50 mM KCl (Ambion #AM9640G), 200 µg/mL CHX (Thermo, J66004.XF), 1 mM DTT (Invitrogen #707265ML), 1X cOmplete EDTA-free Protease Inhibitor (Roche #11836170001), RNasIN Plus 8 U/mL (Promega #N2611), 1% Triton X-100 (Sigma #T9284-1L)). Lysis was performed in 2 mL DNA LoBind tubes (Eppendorf #0030108078) using Zirconia/Silica beads (Biospec #11079105z) in a ThermoMixer C (Eppendorf #5382000015) at 4°C with 20 cycles of 30 s ON at 2000 RPM / 30 s OFF. Lysates were subsequently clarified by spinning at 10000g for 10 min at 4°C. RNA concentration in the supernatant was estimated by measuring absorbance at 260 nm in UV-cuvette micro (Brand # 759235). 3.5 A_260_ units were loaded onto 20-50% sucrose gradients prepared with polysome gradient buffer (PGB = PLB with no Triton-X) in Ultraclear Centrifuge Tubes (Beckman Coulter #C14293). Ultracentrifugation was performed using the SW41Ti rotor at 40 000 rpm for 2.5 h at 4°C. We generated polysome profiles using a BioComp Gradient Station machine and collected 25 fractions of 450 µL each. Fractions were stored at −80°C until further use.

Fractions corresponding to the same profile peak (e.g. monosome, disome, etc.) were combined, and spike-in synthetic RNAs were added to each peak. We extracted the samples using an equal volume of 25:24:1 phenol:chloroform:isoamyl alcohol, obtained by mixing equal parts of citrate saturated phenol pH 4.5 (Sigma #P4682-400mL) and 24:1 chloroform:isoamyl alcohol (Sigma #25666-500mL). Glycoblue (Invitrogen #AM9515) was added according to manufacturer’s instructions, and the aqueous phase was precipitated with 0.1 volumes of sodium acetate pH 5.5 (Invitrogen #AM9740) and 2.5 volumes EtOH overnight at −80°C. RNA pellets were recovered by spinning 20 000g for 30 min at 4°C, washed in 75% EtOH, and subsequently resuspended in 40 µL of RNase-free H_2_O. RNAs were then purified with RNA Clean and Concentrator columns (Zymo #R1016) according to manufacturer’s instructions including DNase treatment (Zymo #E1011).

RNA was quantified with RT-qPCR as mentioned in the RNA quantification methods section, with the mRNAs corresponding to each peak being normalised to the spike in intensities. Polysome propensity, fractional ribosome occupancy of mRNAs and mean number of ribosomes associated with an mRNA were calculated as previously described^62^.

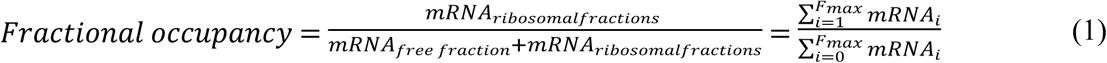

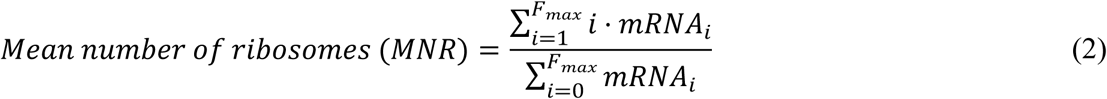

Here, *mRNA_i_* denotes the abundance of a specific mRNA in the *i*-th sucrose gradient fraction, corresponding to the number of ribosomes. Fraction 0 stands for the free mRNA (non-ribosomal) fraction. *F_max_* is the collected fraction with largest number of ribosomes.

### RNA quantification

Reverse transcription was carried out with SuperScript IV (ThermoScientific #18090200), using gene specific primers according to the manufacturer’s instructions. qPCR was then carried out (Roche LightCycler ® 480, KAPA SYBR Fast #KK4611) to obtain Cp values.

For the mRNA decay and translation capacity assessment experiments, qPCR was carried out with gene specific primers. For the translation efficiency calculation, qPCR was carried out with a common primer pair across constructs to reduce amplification-based artefacts.

For mRNA decay and translation efficiency experiments, mRNA levels were normalized to the endogenous housekeeping genes *TFC1* and *UBC6*, unless otherwise stated. For the translation capacity experiments, mRNA levels were normalised to the spike in controls.

To exclude genomic DNA contamination, qPCR was performed on mock reverse transcription reactions (–RT control). For all tested mRNAs spanning a broad expression range, Cp values in –RT samples were near the detection limit (Cp ≈ 35), corresponding to >100-fold and typically >1000-fold lower signal compared to +RT samples (Data S7). These results indicate that residual genomic DNA does not contribute appreciably to the measured mRNA signals.

### Run-off assay

Cells were grown at 30°C in complete synthetic media containing 2% (w/v) glucose until the mid-log phase (OD_600_ = 0.5). To initiate the run-off, cells were filtered and resuspended in glucose-free medium for 2.5 minutes prior to sampling. A control sample (0 min) was collected from the mid-log culture immediately prior to filtration.

At each timepoint, two parallel aliquots were harvested. One aliquot was processed for polysome profiling and RNA quantification as previously described in the Polysome profiling section. The second aliquot was collected in chilled methanol for total mRNA isolation using the RNeasy Mini Kit (Qiagen #74104). To ensure volumetric normalization, spike-in RNAs were added to both total and polysome-fractionated samples.

Purified RNA from both polysome-fractionated and total mRNA samples was quantified by qRT-PCR. Reverse transcription (RT) was carried out with SuperScript IV (Thermo #18090010) using oligo(dT)_15_ (Promega #C1101) and qPCR was then carried out using Roche Lightcycler 480 with KAPA SYBR Fast (Sigma #KK4611). mRNA readouts were normalized to the geometric mean of the spike-ins and then used for further analysis.

The run-off speed was calculated as the ratio of mRNA in the free fraction at 2.5 minutes relative to the 0-minute baseline, with each value normalized to the corresponding total mRNA levels.

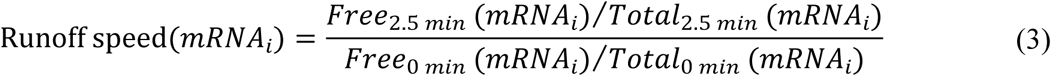

### Systematic comparison of relative elongation speeds determined by runoff and TE-based estimates

To select mRNAs for a systematic comparison, we performed a three-stage selection procedure. First, we established a robust dataset by selecting mRNAs for which unified Elongation Speed (ES) and Translation Efficiency (TE) values were available from all three referenced TE studies and for which the coefficient of variation (CV) across these measurements was CV<1. This initial filtering resulted in 491 mRNAs.

Second, we employed a 2×2 stratified sampling design to identify mRNAs representing the range of CDS lengths where elongation speed is observed to vary as a function of codon optimality. This design crossed CDS length (short vs. intermediate) with the mean codon stabilization coefficient (μ_CSC_; low vs. high).

µ_CSC_ categories were defined using the bins shown in Fig. 4H: “high” corresponded to the top bin, µ_CSC_ ∈ (0.2, 0.366], and “low” to the bottom bin, µ_CSC_ ∈ [−0.106, 0]. Within each µCSC category, genes were drawn from two CDS-length windows centered on 600 nt (“short”) and 1200 nt (“intermediate”). Formally, the short set was defined as S = {g ∈ G : |L(g) − 600 nt| ≤ ε} and the intermediate set as I = {g ∈ G : |L(g) − 1200 nt| ≤ ε}, with ε grown stepwise from zero until each set contained exactly four genes. The final ε was < 100 nt in all four categories.

Third, runoff speeds were measured for these 16 candidate mRNAs. We then performed a final quality control step by selecting only those mRNAs whose relative elongation speed, as determined by the runoff, had a CV<0.5. This multi-stage procedure resulted in a final set of 13 mRNAs, representing a highly reliable dataset for both the runoff-derived and TE-based variables, thereby providing a solid basis for correlation analysis with low measurement noise.

### Generation of Variable Poly(A) Length Transcripts

To synthesize mRNA constructs with specific poly(A) tail lengths, we adapted a PCR-based strategy from Grandi et al^63^. Briefly, eGFP-mScarlet was amplified templated by pSYN129 using Q5 Hot Start High-Fidelity DNA Polymerase (NEB, #M0493S). Forward primers were designed to introduce a T7 promoter sequence, optimal spacer sequences, and length-specific 9-nucleotide barcodes for downstream qPCR selectivity. Reverse primers were designed with 5’ poly(T) tracts of desired poly(A) tail lengths (0 to 75 nt).

Amplicons were verified by 1% agarose gel electrophoresis and Sanger sequencing, then purified using the NucleoSpin Gel and PCR Clean-up kit (Macherey-Nagel, #740609.250).

In vitro transcription was performed using 1 µg of purified template DNA and the TranscriptAid T7 High Yield Transcription Kit (Thermo Scientific, #K0441) with an incubation at 37°C for 4 hours. Template DNA was subsequently degraded using 2 U of DNase I (Zymo Research, #E1009-A) at 37°C for 15 minutes. RNA transcripts were purified via acidic phenol-chloroform (pH 4.5) extraction, followed by overnight precipitation at −80°C with 0.1 volume of sodium acetate, pH 5.5 (Thermo #AM9740), GlycoBlue (Thermo # AM9515), and three volumes of 100% ethanol. RNA pellets were collected by centrifugation (20,000 × g, 1 h, 4°C), washed with ice-cold 75% ethanol, air-dried, and resuspended in nuclease-free water. RNA was quantified and diluted to 250 ng/µL for storage at −80°C.

For downstream quantification, qPCR primers were designed to specifically target the unique 9-nt barcode sequences associated with each poly(A) construct.

### ePAT

Yeast strains harbouring constructs with silent mutations to enable multiplexed gene control experiments were cultured and harvested in mid-log phase. The ePAT protocol was performed with modifications to the method described by Jänicke et al.^64^ in order to enable the detection of genes containing silent mutations in the 5’ region of their coding sequences (CDS)

In the first stage, the poly(A) tail extension and cDNA synthesis were performed. 500 ng of RNA (total or poly(A)-selected, with yeast tRNA added as ballast for the poly(A)-selected samples) was annealed to 1 µL of 100 µM PAT anchor primer at 80 °C for 5 min. 3’ ends were then extended using 5 U of DNA Polymerase I, Large (Klenow) Fragment (NEB #M0210M) in 1× SuperScript IV buffer (Invitrogen #18090050) supplemented with dNTPs, DTT and RNaseIN (Promega #N2615) at 37 °C for 1 hr. Klenow was inactivated at 80 °C for 10 min, the reaction was brought to 55 °C, and 1 µL of SuperScript IV Reverse Transcriptase (200 U; Invitrogen) was added directly to the heated tubes; reverse transcription proceeded for 1 hr at 55 °C followed by a final 80 °C inactivation.

In the second stage, the primary PCR was carried out to enrich the target templates. The resulting cDNA was diluted 1:10 and subjected to a 10-cycle primary PCR using Q5 Hot Start High-Fidelity DNA Polymerase (NEB #M0493S) with the PAT anchor reverse primer and gene-specific forward primers that recognizes the silent mutations. Cycling conditions were: 98 °C for 1 min; 10 cycles of 98 °C for 10 s, 66 °C for 30 s and 72 °C for 30 s per kb of amplicon; and a final extension at 72 °C for 5 min.

In the third stage, a semi-nested secondary PCR was performed to increase specificity and add a GC-clamp for resolution. The primary PCR product was diluted 1:500 and used as template for a subsequent PCR with GoTaq G2 Master Mix (Promega #M7822), the PAT GC-clamp reverse primer, and 3’ UTR-specific forward primers. Cycling conditions were: 95 °C for 2 min; 31 cycles of 95 °C for 15 s, 56 °C for 15 s and 72 °C for 30 s; and a final extension at 72 °C for 5 min.

In the fourth stage, the resulting PCR products were resolved and analyzed. PCR products from stage 3 were resolved on a 4% low-range agarose gel (Bio-Rad #1613106) prestained with RedSafe (Luzerna Chem #21141) at 70 V for 150 min, imaged, and analyzed using Fiji.

### ePAT–qPCR assay

The ePAT–qPCR assay extends the previously described ePAT method to enable selective quantification of short poly(A)-tailed (∼A12) transcripts using qPCR as the readout. The assay proceeds in three stages (Fig. S5F). (1) Klenow extension and reverse transcription of poly(A)-tailed mRNA using a TVN-anchored PAT primer, (2) a 1st round PCR that amplifies target transcripts from the expressed constructs via a barcode-containing forward primer, and (3) a qPCR stage (2nd round PCR) in which a tripartite reverse primer selectively reports on transcripts carrying short (∼A12) tails (see primer sequences in Data S6). Because the qPCR reverse primer contains a fixed T12 stretch flanked by UTR-complementary sequence and the PAT anchor, it hybridizes efficiently only when the poly(A) tail is short enough that the UTR and anchor sequences sit in close proximity, thereby providing a readout of only the short poly(A)-tailed species.

Yeast cells were cultured, collected, and RNA was extracted as described in the ePAT section. Where indicated, 50 µg of total RNA was subjected to poly(A) selection using the PolyATtract mRNA Isolation System IV (Promega #Z5310) following the manufacturer’s protocol, with final elution in 250 µL.

In the first stage, the poly(A) tail extension and cDNA synthesis were performed to anchor the mRNA 3’ ends. 1 µg of total RNA—or 5 µL of poly(A)-purified material supplemented with yeast tRNA to a total mass of 1 µg—was combined with 1 µL of 100 µM TVN PAT anchor primer and subjected to Klenow extension and reverse transcription as described in the ePAT section.

In the second stage, the 1st round PCR was carried out to enrich the target templates containing the silent barcode mutations. The resultant cDNA from Stage 1 was diluted 1:10 with nuclease-free water. 5 µL of diluted cDNA was amplified in a 25 µL reaction using Q5 Hot Start polymerase (NEB #M0493S) with reagents at the manufacturer’s recommended concentrations. The forward primer targeted a region within the coding sequence containing four silent barcode mutations that distinguish the TET-off construct from the endogenous gene. The TVN PAT anchor primer served as the reverse primer. Endogenous TFC1 and UBC6 served as internal controls, with forward primers designed to yield amplicons of similar size. Cycling conditions were: 98 °C for 30 s (initial denaturation); 15 cycles of 98 °C for 30 s, 65 °C for 30 s, 72 °C for 1 min; and a final extension at 72 °C for 5 min.

In the third stage, the 2nd round PCR (qPCR) was performed using a tripartite primer to quantify the proportion of short-tailed transcripts. The 1st round PCR products were diluted 1:1000 with nuclease-free water. 2 µL of this dilution was used in a 10 µL qPCR reaction with 5 µL 2x KAPA SYBR Fast (Sigma #KK4611) and 0.1 µM of forward and reverse primers. The forward primer was placed in the 3ʹ UTR to yield an amplicon of 75–150 nt. The reverse primer was designed as a tripartite oligonucleotide (5ʹ→3ʹ): 5–7 nt of the PAT GC clamp, a T12 stretch, and 5–7 nt complementary to the 3ʹ UTR immediately upstream of the polyadenylation site. This design ensures efficient amplification only when the poly(A) tail is sufficiently short (∼12 nucleotides) that both the UTR-complementary and anchor-complementary segments of the primer engage their respective templates, thereby selectively reporting on short-tailed species of a defined UTR isoform. Since most mRNAs have several 3’ UTR isoforms^65^, multiple isoforms per mRNA were probed such that they together comprised >75% of the total mRNA abundance (Table SX). qPCR reactions were run on a Roche LightCycler 480.

For each major 3’UTR isoform (*i*) associated with an mRNA (*m*), the relative abundance of the short poly(A)-tailed transcript was measured using a forward primer recognizing the UTR and the tripartite reverse primer specific for both a given UTR isoform and the (A)_12_ tail. The total mRNA was measured with primers annealing to the silent mutations in the CDS, ensuring a readout of the entire population regardless of tail length or UTR isoform. The relative abundance of short poly(A)-tailed species for each isoform was calculated by dividing the isoform-specific short-tailed level by the total mRNA level, [*m*]_i_/ [*m*]_total_. The short poly(A)-tailed proportion for each mRNA was then calculated by summing these ratios across all measured UTR isoforms:

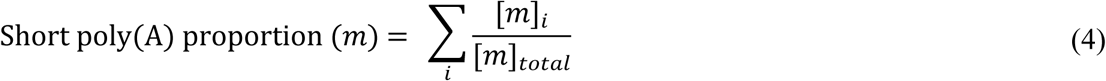

### Protein Quantification by ELISA

Cells were collected by spinning down 5mL of the culture by centrifugation at a speed equivalent to 2000g, at a temperature of 4°C. The supernatant was removed and the cell pellet resuspended in 1x Cell Extraction Buffer PTR (from GFP SimpleStep ELISA® Kit, Abcam #ab171581). Cell suspensions were loaded into 1.5-ml microfuge tubes on benchtop shaker/incubator set at 4°C, and lysis was carried out with Zirconia/Silica beads (Biospec #11079105z). Twenty cycles of interval shaking were carried out with each cycle consisting of 30s shaking at 2000rpm and 30s resting periods. The tubes were then spun at 10,000g for 5min to clarify the lysate.

Total protein quantification was carried out on the lysates using bicinchonic acid (BCA) assay (Pierce™ BCA Protein Assay Kit, ThermoFisher Scientific #23225), using 1x Cell Extraction Buffer PTR as the diluent for the standards. Measurements were carried out in microplate format (Greiner #655160), on a Tecan Spark® Multimode plate reader at 562nm. Known quantities of total protein (ranging between 5ng and 50ng, so as to be within the calibration range of the assay) were then used for ELISA against GFP in a sandwich ELISA strategy. The assay was carried out as per kit protocol (GFP SimpleStep ELISA® Kit, Abcam #ab171581). Measurements were carried out in an endpoint reading manner, with the stop solution being added 10min after addition of the TMB Development Solution. The samples were measured on a Tecan Spark® Multimode plate reader at 450nm and GFP was quantified using a standard curve with a quadratic fit as described in the protocol.

### Protein Quantification by Flow Cytometry

Samples were collected in 5 mL round-bottom polystyrene tubes (BD Falcon #352054) and maintained on ice. Data acquisition was performed using a BD LSR Fortessa™. Single-cell populations were isolated via standard FSC and SSC gating. Protein expression was quantified using mean fluorescence intensity for GFP-A (488 nm laser, 505LP, 512/25 filter) and mScarlet-A (561 nm laser, 600LP, 610/20 filter).

### Determination of protein half-lives

To measure the degradation kinetics of GFP-RFP fusion proteins, cultures were grown overnight at 30°C to an OD600 of 0.5–1.0. Expression was repressed by adding doxycycline (Sigma #D9891) to a final concentration of 10 µg/mL. Samples were collected at 0, 2, 4, 6 and 8 hours post-treatment; the 0-hour sample was taken immediately prior to doxycycline addition. Logarithmic growth was maintained throughout the assay by diluting cultures 1:2 with fresh media every two hours.

Time-course mean fluorescence intensity data for each strain were fit to a single-exponential decay model: ·𝑦 = A 𝑒^―𝑘⋅𝑡^

Fits were performed via nonlinear least-squares regression (scipy.optimize.curve_fit). Standard errors for A and k were obtained from the covariance matrix of the fit, and the half-life was computed as t½ = ln(2)/k with its standard error propagated via the delta method. Goodness of fit was assessed by R² and adjusted R².

To test whether the strains express proteins of differing half-life, an omnibus extra sum-of-squares F-test was first performed, comparing an unconstrained model in which each sample had its own decay rate to a constrained model in which all samples shared a single common rate (with each sample retaining its own amplitude).

Normalization of translation efficiency datasets

To analyze translation efficiency (TE), three datasets reporting translation rate constants (see section Data analysis of Translation Efficiency and Elongation speed) were collected^17–19^. The study by Riba et al. contained two datasets that differed by the mRNA isolation: one measured with poly(A) selection in the same study (Riba dataset) and one measured after rRNA depletion in a separate study (Riba RiboZero dataset)^26^.

Four approaches were evaluated for normalizing the datasets before unification. In all cases, only genes represented in all three datasets were considered, and the coefficient of variation (CV) across datasets was calculated to assess measurement error.

**1. Unit conversion.** Datasets were converted to the same units without additional normalization.
**2. Median normalization**. Values in each dataset were normalized to the dataset median.
**3. Regression-based normalization.** Translation efficiency was predicted for each mRNA from its average codon stabilization coefficient (µ_CSC_) using linear regression (Fig. S1B). The median of predicted TE values was then used for normalization instead of the median of measured TEs, thereby reducing both measurement errors and the insufficiency caused by the limited number of shared genes used in standard median-based normalization. Regression variables were calculated from the common gene set within each dataset.
**4. Scaling factor normalization.** Scaling factors were freely varied and selected to minimize the average CV across datasets.

The performance of each approach was compared. Median normalization (approach 2) is generally robust^16,40^, but in this case simple unit conversion (approach 1) yielded lower CVs. Regression-based normalization (approach 3) reduced measurement errors and mitigated problems arising from the limited overlap in gene coverage, and also improved performance for mRNAs with long coding sequences (>1,000 nt), but overall performed similarly to simple unit conversion. Scaling factor normalization (approach 4) yielded a slightly lower CV than regression-based normalization, but again did not outperform simple averaging. On this basis, the simplest approach (unit conversion without normalization) was selected for dataset unification.

### Unification of Translation Efficiency datasets

A unified TE value was calculated for genes that showed consistent TE estimates across multiple datasets. For each gene with TE values available in more than one dataset, the individual TE values were arranged in ascending order 𝑥_1_,𝑥_2_,𝑥_3_where the total number of available datasets is denoted as |𝑋| = 𝑛, and dataset-specific percentiles are denoted as P_i_.

When only one dataset provided a TE value for a given mRNA (|𝑋| = 1), this value was accepted if it was not considered an outlier—specifically, if it fell between the 2.5th and 97.5th percentiles of TE values within that dataset:

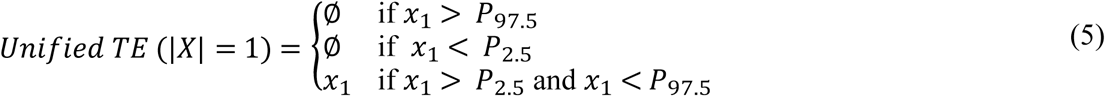

When two datasets provided TE values (|𝑋| = 2), the unified TE was calculated as the geometric mean of the two, provided that their values were within one order of magnitude (i.e., < 10-fold difference). If the values differed by more than 10-fold, no unified TE was assigned:

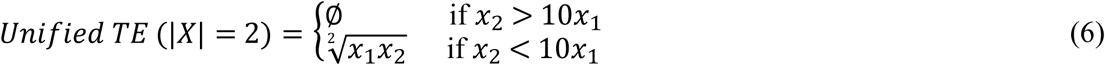

For genes with TE values in three datasets (|𝑋| = 3), the unified TE value was determined based on the pairwise ratios of the ordered TE values: 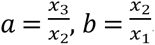.

If both ratios, *a* and *b* exceed 10, no unified TE value was assigned.

If the overall ratio between the highest and lowest TE values (𝑎𝑏 = 𝑥_3_/𝑥_2_ ) was less than 10, the unified TE was computed as the geometric mean of all three TE values. If the total spread exceeded 10-fold, but only one of the pairwise ratios (*a* or *b*) was large, the unified TE was calculated using the two most consistent values (those differing by ≤10-fold).

In cases where both 𝑎 and 𝑏 were ≤ 10 and equal in value, yet the ratio between the highest and lowest TE values exceeded 10, no unified TE value was assigned, because such a disparity also indicates inconsistent TE range for the mRNA.

Formally:

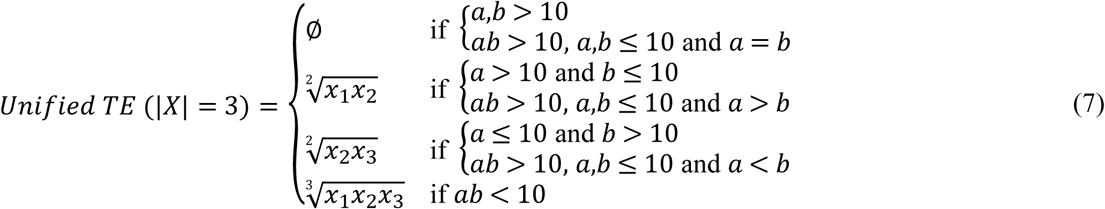

### Extrapolation of the unified TE dataset to total RNA

To extrapolate the unified TE values from poly(A)+ to total RNA, we adjusted them using data from two studies: Riba et al., which provided paired mRNA abundance measurements for poly(A)+ and total RNA (RiboZero) samples, and Presnyak et al., which reported corresponding mRNA half-lives (t₁/₂) for both RNA types.

A linear regression model was established to predict the ratio of poly(A) to total mRNA abundance, using the ratio of poly(A) to total mRNA half-life as the predictor variable. The model’s parameters, 𝑎 and 𝑏, were fitted using data from 737 genes common to both the Riba and Presnyak datasets:

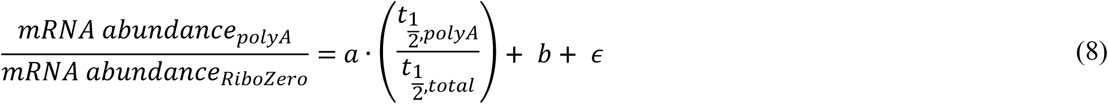

This model was then used to predict the mRNA abundance ratio for 2,138 genes that had both a unified TE value, derived from poly(A)-selected RNA datasets, and decay rate measurements in the Presnyak dataset (Data S4). The extrapolated TE, representative of total RNA levels, was calculated by multiplying the initial poly(A)-based unified TE by the predicted abundance ratio:

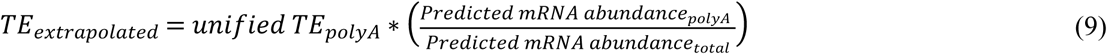

As TE is inversely proportional to mRNA abundance, this multiplication adjusts the initial value by canceling the poly(A)-selection bias, yielding a TE estimate that reflects translational efficiency relative to the total cellular mRNA pool.

### Data analysis of Translation Efficiency and Elongation speed

In this study, TE is defined as the rate constant for translation, *k_t_*, according to the differential equation that describes translation of mRNAs into proteins:

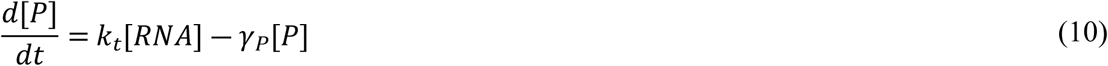

where P denotes the protein concentration, γthe protein decay rate constant, and [RNA] the mRNA concentration.

In the unified dataset, relying on the proteomic measurements, *k_t_* was estimated directly from the measured protein abundance and decay rate (determined by SILAC), together with mRNA abundance.

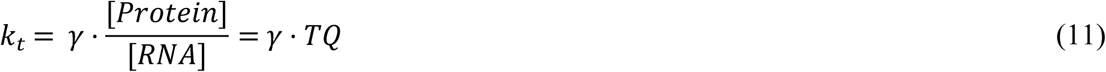

The translation quotient (TQ) corresponds to the ratio of the steady-state protein to RNA abundance. For the synthetic mRNAs, the relative translational efficiency (Rel. TE) was estimated as the ratio of 𝑘_𝑡_of the mRNA of interest (ROI) relative to a reference mRNA;

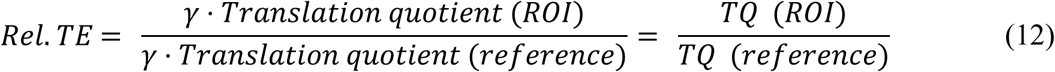

In the TQ, the steady-state abundances of the protein and RNA are normalized either to total or specific species. Because all synthetic constructs encode the same protein, they were assumed to share the same protein decay rate constant.

For calculating the relative TE with respect to poly(A)⁺ mRNA, the reference strain’s value (set to 1) was adjusted by the poly(A)⁺/total mRNA ratio for each replicate experiment.

From the TE values, we derived additional parameters, including the codon translation coefficient (CTC) and the elongation speed.

CTC is the Pearson correlation coefficient between the frequency of a codon across mRNAs and the corresponding translation efficiencies. The related metric for mRNA stability, the CSC values were obtained from a study using multiplexed gene control to measure mRNA half-lives at 30°C^7^.

To relate these measures to tRNA availability and codon bias, data on tRNA gene copy number (tGCN), microarray-based, tRNA-seq, and OTTER-based tRNA abundances were obtained from Nagai et al^20^. Codon adaptation index (CAI), the tAI and its variants (stAI and gtAI) were obtained from Anwar et al^66^.

While tAI was originally parameterized using S. cerevisiae gene expression data^31^, the species-specific tAI (stAI) has been shown to underperform the original tAI in predicting protein abundance across fungal genomes. We therefore also included the genetic tAI (gtAI), which uses a genetic-algorithm-based optimization to address limitations of stAI’s hill-climbing approach.

Elongation speed was inferred from TE using the continuity equation^27^. Assuming a steady-state ribosome distribution along the mRNA, the flux of ribosomes through codon x is given by: 𝐽(𝑥) = 𝑣(𝑥)𝜌(𝑥)

where *v(x)* is the translation speed at position *x*, and *ρ(x)* is the ribosome density at that position. The ribosomal current J corresponds to the protein production rate per mRNA (i.e., the number of proteins produced per unit time per mRNA)^61^.

For an average velocity,

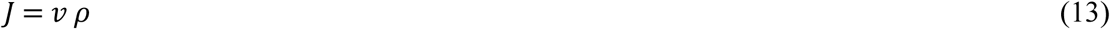

This formulation can be viewed as an approximation for translation on a ring, which closely describes ribosomal reinitiation following a mRNA circularization between the stop and start codons^67^, rather than the open-end approximations typically used for translation^68^.

Modelling of degradation of total and poly(A)+ mRNA

The shortening of poly(A) mRNA occurs in multiple steps. These equations are formulated according the observation that poly(A) shortening and mRNA decapping are not strictly coupled and decapping and the subsequent 5’-to-3’ degradation can occur even in the presence of poly(A) tails^11^. For simplicity, four major species are distinguished according to poly(A) length: long (L), medium (M), short (S), and null (N). Each of these is shortened progressively at rates *k_da1_, k_da2_* and *k_da3_,* , resulting in the conversion of each mRNA species into the next shorter species. Once the poly(A) tail is completely de-adenylated, the mRNA can be degraded in the 3’-to-5’ direction be the exosome at a rate of 𝛾_𝐸𝑥𝑜_.

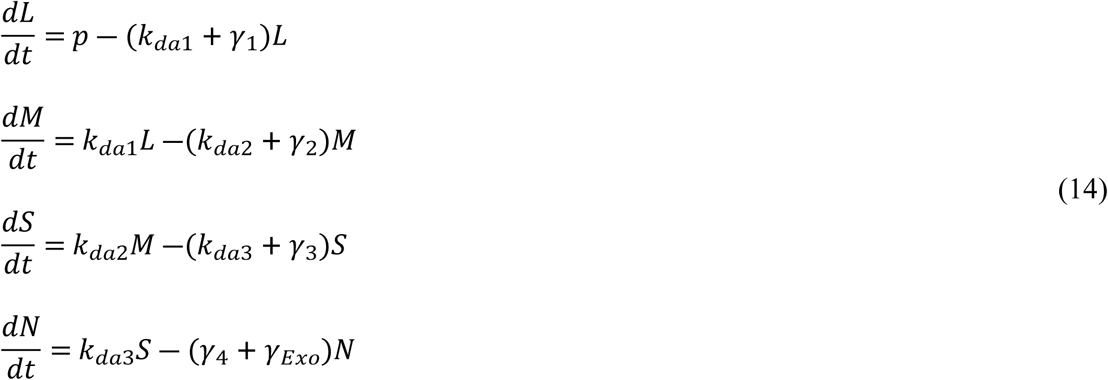

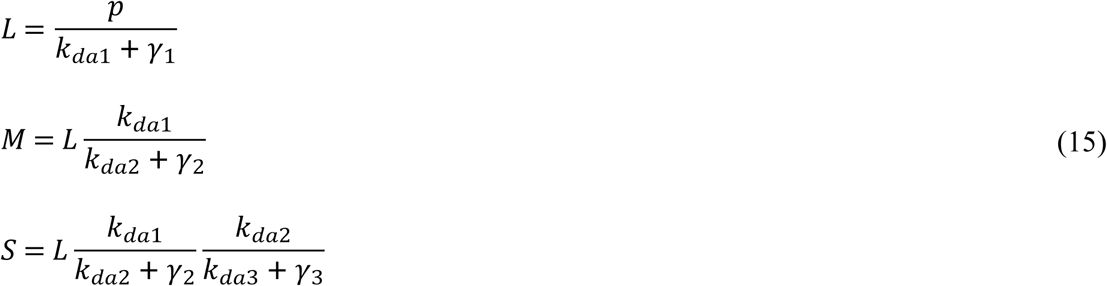

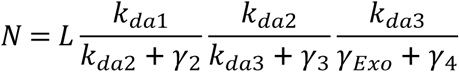

Assuming that the decay rates of the mRNAs with poly(A) tails of different lengths are equal, the following simplification can be done since all mRNAs with a poly(A) tail are degraded by Xrn1 in the 5’-to-3’ direction: 𝛾_1_ = 𝛾_2_ = 𝛾_3_ = 𝛾_4_ = 𝛾_𝑋𝑟𝑛1_

The deadenylation rates can differ according to poly(A) length^13^, and it is highest for intermediate length.

The steady-state levels are accordingly

The following relations were established to associate the mRNA species with the appropriate pool of mRNA isolated with a specific technique:

Total mRNA = N + S + M + L

mRNA isolated with LNA beads = S + M + L

mRNA isolated with Oligo-dT beads = M + L

### Outlier removal in the Half-life dataset obtained by transcriptional inhibition

Outlier removal for the transcriptional inhibition dataset (Presnyak) was performed following the unification procedures described previously^4,16^. Multiple half-life (t1/2) datasets were unified, and the values in each set were normalized by their respective median half-life. To ensure the reliability of the measurements for each transcript, a trimming procedure using the Median Absolute Deviation (MAD) metric was employed to omit outliers. The median and MAD were calculated for each mRNA across its available measurements in the unified dataset. Half-lives were retained only if they obeyed the following relation:

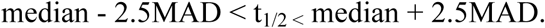

Values falling outside this range were removed. In this study, the Presnyak dataset following this outlier removal is denoted as “Presnyak (outlier-filtered)”.

### Bayesian Estimation of Codon-Specific Elongation Speed

Codon-specific elongation speeds were estimated using Bayesian linear regression. Average gene-level elongation speeds, derived from translation efficiency (TE) and ribosomal density, were combined with codon frequencies of each gene to infer codon-level contributions.

The regression model included 61 variables corresponding to the 61 sense codons:

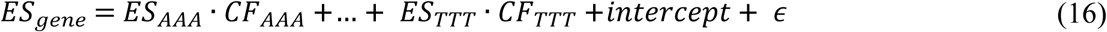

where *ES* denotes elongation speed and *CF* the codon frequencies. Stop codons (TAA, TAG, TGA) were excluded.

To incorporate the biological constraint that codon-specific elongation speeds must be positive, a Gamma prior distribution was imposed:

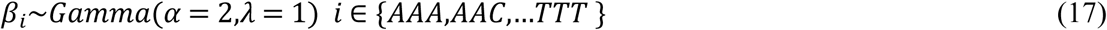

The model was implemented with the **brms** package in R^69^. Priors were set using the set prior function, and models were fit using brm. Posterior estimates of codon-specific elongation speeds, along with confidence intervals and error estimates, were obtained for different gene subsets (Data S3).

### Calculation of Control Coefficients

The effects of codon optimality on mRNA half-lives, TE, and elongation speeds were quantified by performing robust linear regression against, the tRNA adaptation index (tAI), which quantifies codon optimality independently of these output variables. Normalization of the output variables enables comparison across distinct quantities and allows the resulting slope to be interpreted as a control coefficient. Because the normalization is discrete rather than continuous, this coefficient differs from control coefficients based on logarithmic derivatives^70^.

The regression was performed with the Theil–Sen estimator, which determines the slope of the relationship between tAI and the output of interest by computing slopes for all pairs of data points and taking the median. Confidence intervals for the slopes were computed following Sen’s method^71^, using the using ci.theil command in statpsych (v1.8.0) [Bonett D (2025).

statpsych: Statistical Methods for Psychologists. R package version 1.8.0, https://dgbonett.github.io/statpsych/] package of R (v4.5.1).

### Calculation of P-values with permutation test

Two-tailed P-values were estimated by Monte Carlo sampling using^72^:

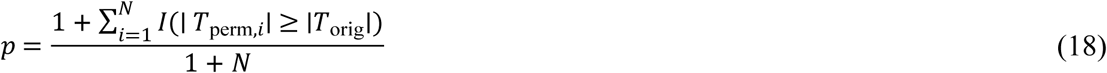

where N is the number of permutations, Tₒᵣᵢg is the test statistic computed from the observed data, Tₑᵣ,ᵢ is the test statistic recomputed from the ith permutation, and I() denotes the indicator function. The pseudo-count of 1 in numerator and denominator guarantees a minimum attainable P-value of 1/(N + 1). All tests were two-tailed. Unless otherwise stated, N = 10,000, yielding a minimum P-value of ≈ 10⁻⁴.

The null hypothesis, the test statistic T, and the resampling scheme employed in each test are described below.

For the differences between control coefficients, the null hypothesis was that the control coefficients for two output variables (e.g. half-life and TE) are equal. The test statistic was the difference between the corresponding Theil–Sen slopes [ΔTS = TS(t₁/₂) − TS(TE)]. For each permutation, half-life and TE values were randomly shuffled across genes, and ΔTS was recalculated.

For the pairwise correlations between parameters, the null hypothesis was that the two parameters are statistically independent. The test statistic was the Spearman correlation coefficient ρ between them. For each permutation, the values of one parameter were randomly shuffled across genes or codons, and ρ was recalculated.

For the CSC and CTC, the null hypothesis was that the specific arrangement of codons within a gene does not affect its half-life or TE beyond what is expected by chance for a sequence of that length and composition. The test statistic was the CSC or CTC value of a given codon. For each permutation, the codon frequencies were randomly shuffled among the codons within each gene, and thereby breaking the structural dependencies that arise because codon frequencies are compositional (i.e., the 64 codons are analyzed as a closed set)—and the test statistic was recalculated. N = 10,000 for CTC-based tests and N = 100,000 for CSC-based tests (minimum P ≈ 10⁻⁵).

